# A natural fusion of flavodiiron, rubredoxin, and NADH:rubredoxin oxidoreductase domains is the highly efficient water-forming oxidase of *T. vaginalis*

**DOI:** 10.1101/2022.02.02.478753

**Authors:** Evana N. Abdulaziz, Tristan A. Bell, Bazlur Rashid, Mina L. Heacock, Owen S. Skinner, Mohammad A. Yaseen, Luke H. Chao, Vamsi K. Mootha, Antonio J. Pierik, Valentin Cracan

## Abstract

Microaerophilic pathogens such as *Giardia lamblia* and *Trichomonas vaginalis* have robust oxygen consumption systems to detoxify oxygen and maintain the intracellular redox balance. This oxygen consumption is a result of the H_2_O-forming NADH oxidase activity of two distinct flavin-containing systems: H_2_O-forming NADH oxidases (NOXes) and multicomponent flavodiiron proteins (FDPs). Both systems are not membrane-bound and recycle NADH into oxidized NAD^+^ while simultaneously removing O_2_ from the local environment, making them crucial for the survival of human microaerophilic pathogens. In this study, using bioinformatic and biochemical analysis, we show that *T. vaginalis* lacks a NOX-like enzyme, and instead harbors three proteins that are very close in their amino acid sequence and represent a natural fusion between N-terminal FDP, central rubredoxin and C-terminal NADH:rubredoxin oxidoreductase domains. We demonstrate that this natural fusion protein with fully populated flavin redox centers unlike a “stand-alone” FDP (also present in *T. vaginalis*), directly accepts reducing equivalents of NADH to catalyze the four-electron reduction of O_2_ to water within a single polypeptide and with an extremely high turnover. Using single particle electron cryo-microscopy (cryo-EM) we present structural insight into the spatial organization of the FDP core within this multidomain fusion protein. Our studies represent an important addition to our understanding of systems that allow human protozoan parasites to maintain their optimal redox balance and survive transient exposure to oxic conditions.

## Introduction

*Trichomonas vaginalis* is a microaerophilic human protozoan parasite that causes trichomoniasis, one of the most common sexually transmitted infections (1,2). A distinct feature of *T. vaginalis* as well as other human parasites such as *Giardia lamblia* (syn. *intestinalis, duodenalis*) and *Entamoeba histolytica* is their ability to tolerate low oxygen concentrations and transient exposure to oxic conditions. This is remarkable as these protozoa depend on metabolic enzymes that are extremely sensitive to oxygen, such as pyruvate:ferredoxin oxidoreductase (PFOR) and FeFe-hydrogenase (3,4). These organisms do not contain mitochondria, and instead harbor mitochondria-derived organelles called mitosomes or hydrogenosomes that lack a membrane-bound electron transport chain (ETC) to carry out oxidative phosphorylation (OXPHOS) (4). Moreover, it is well recognized that *T. vaginalis* consumes oxygen and that this oxygen consumption is insensitive to ETC inhibitors (such as cyanide or sodium azide) (4,5). Over the years, most of this oxygen consumption was attributed to enzymes that catalyze a conversion of diatomic oxygen to two benign water molecules at the expense of reducing equivalents of NAD(P)H (the H_2_O-forming NAD(P)H oxidase reaction) (**Fig. 1A**). This reaction not only serves the “oxygen scrubbing” role but at the same time maintains uninterrupted supply of oxidized NAD^+^ to support optimal intracellular NADH/NAD^+^ ratio (4,6).

**Figure 1.**
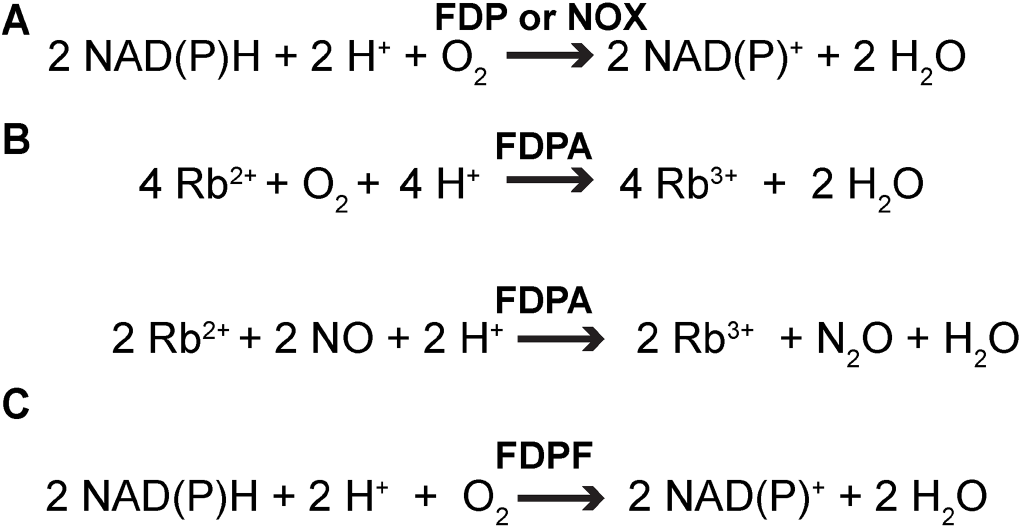
Reactions catalyzed by Class A and Class F Flavodiiron Proteins (FDPA and FDPF) as well as by a H_2_O-forming NADH oxidase (NOX). **(a)** The net H_2_O-forming NAD(P)H oxidase reaction catalyzed by a multicomponent flavodiiron protein (FDP)-based system or a H_2_O-forming NADH oxidase (NOX). **(b)** In the simplest configuration, Class A FDP (FDPA) receives electrons from reduced rubredoxin (Rb). Some FDPAs can also reduce nitric oxide to nitrous oxide and water, respectively. (**c**) Class F FDP (FDPF) represents a natural fusion, where the FDP core protein is fused with both rubredoxin (Rb) and NADH:rubredoxin oxidoreductase (NROR) redox partners.

Through convergent evolution, two different enzymatic systems evolved to catalyze the H_2_O-forming NADH oxidase reaction in various bacteria and protozoa (**Fig. 1A**). The first system is an H_2_O-forming **<u>N</u>**ADH **<u>ox</u>**idase (NOX) that belongs to the evolutionarily versatile “two dinucleotide binding domains” flavoproteins (tDBDF) superfamily (**Fig. 1A**) (7–9). These soluble NOXes are not related to the mammalian transmembrane ROS-producing NADPH oxidases (NOX1-5 and DUOX1-2). The active site of a typical tDBDF superfamily NOX enzyme (~50 kDa per monomer) consists of a single FAD cofactor and an adjacent redox-active cysteine that cycles between sulfenic acid and reduced cysteine to accomplish the four electron reduction of oxygen (7,10,11). For example, in our previous study we extensively both biochemically and structurally characterized the water-forming oxidase reaction of *L. brevis* NOX (*Lb*NOX) (11). *Lb*NOX is an extremely efficient enzyme with strict specificity towards NADH, with *K*_Ms_ for O_2_ and NADH, of ~2 μM and 69 ± 3 μM, respectively, and the turnover number of 648 ± 28 s^−1^ while less than 2 % of input electrons lead to the off-target H_2_O_2_ formation (11).

Flavodiiron proteins (FDPs) are a second system with H_2_O-forming oxidase activity (**Fig. 1A-C**, **Fig. 2A**). Interestingly, some FDPs can also use NO gas as a substrate or can have mixed O_2_/NO specificity (**Fig. 1B**) (12–15). The minimal unit of a typical FDP (in the new classification Class A FDP, see below) is a ~50 kDa protein with two subdomains: an N-terminal metallo-β-lactamase-like subdomain containing the dinuclear Fe-Fe center (diiron center); and a C-terminal subdomain, containing the FMN redox cofactor (15–17)(**Fig. 2A**). The genomes of human protozoan parasites *T. vaginalis, G. lamblia* and *E. histolytica* encode biochemically verified Class A FDPs (18–20). Within a monomer the distance between the oxygen-binding diiron center and the FMN plane is approximately 40 Å in all reported structures of FDPs, much greater than the ~15 Å maximum distance that allows efficient electron transfer (21). As such, the functional unit of a “stand-alone” Class A FDP is comprised of a “head-to-tail” homodimer that positions the diiron center of one protomer adjacent to the FMN bound of its neighboring protomer (15,17,19).

**Figure 2.**
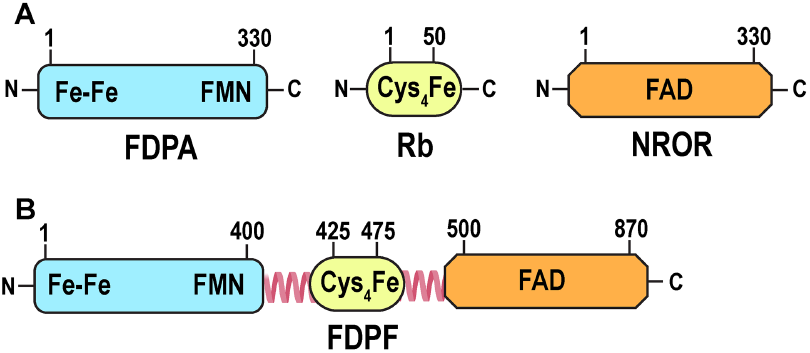
Domain organization of Class A and Class F flavodiiron proteins (FDPA and FDPF). **(a)** Class A flavodiiron protein (FDPA) contains the diiron center (Fe-Fe) and a flavin cofactor (FMN). Two additional redox partners of a “stand-alone” FDPA are: (*i*) rubredoxin (Rb) and (*ii*) NADH:rubredoxin oxidoreductase (NROR) (see Fig. 1). The Fe(SCys)_4_ center of Rb is formed by a central iron which is coordinated by four cysteines. The redox center of NROR contains a flavin redox cofactor (FAD). (**b**) Class F FDP (FDPF) represents a natural fusion between the N-terminal FDP core, Rb-like domain, and NROR-like domain. Loops connecting the middle rubredoxin-like domain with the N-terminal FDP core and the NROR domain are shown.

The FMN redox cofactor of FDP cannot directly receive electrons from NAD(P)H and requires additional protein adaptors as substrates. For example, FDPs can directly receive electrons from a rubredoxin (Rb), a ~ 6 kDa protein that contains a Fe(SCys)_4_ center and in some bacteria and *Archaea* is reduced by NAD(P)H:rubredoxin oxidoreductases (NRORs) or similar systems (4,22–26)(**Fig. 1B**). Notably, in biochemical and spectroscopic studies of previously characterized Class A FDPs, only artificial protein substrates were used (*i.e*. recombinant NADH:flavorubredoxin oxidoreductase and a truncated rubredoxin domain of flavorubredoxin, both from *E. coli*) as in the corresponding organisms Rb and/or NROR homologs were not identified (16,27,28).

Recent bioinformatic analysis revealed that in many organisms FDPs are naturally fused to various domains, and led to the current classification of this large group of proteins (27–29). According to this classification “stand-alone” FDPs that contain only diiron and FMN centers are designated as a Class A FDPs or FDPAs (27) (**Fig. 2A**). Several additional classes were proposed for proteins in which one or more other domains are fused genetically to the C-terminal side of the FDP core (Classes B to H) (27). These additional redox centers may play crucial roles in dedicated electron transfer (ET) pathways, via which reducing equivalents of NAD(P)H or other substrates are relayed to the gas-binding diiron site (27).

The first attempt to purify the H_2_O-forming NADH oxidase activity from *T. vaginalis* isolated a preparation that was active only with NADH (no activity with NADPH was detected) with a specific activity of 16.5 μmol min^−1^ mg^−1^ and *K*_M_ for NADH of 7.4 μM (30). The authors reported that the preparation consumed NADH and O_2_ with 2:1 stoichiometry and did not produce H_2_O_2_ as a byproduct. Unfortunately, neither the N-terminal sequence nor the molecular mass of the enzyme was reported at that time. In a later study, a H_2_O-forming NADH oxidase activity from *T. vaginalis* was purified, and exhibited a V_max_ of 470 s^−1^ and *K*_M_ for NADH of 5.4 μM (31). The authors reported that when analyzed by SDS-PAGE, the final protein preparation migrated as two closely spaced unusually high molecular mass bands of 97-97.5 kDa. The latter observation is inconsistent with NOXes being ~50 kDa proteins (see above) (30–33).

In our studies we turned our attention to the NOX-like activity ascribed to *T. vaginalis* because we were interested in oxygen metabolizing systems that allow unicellular organisms that lack ETC to maintain an optimal redox environment (11). To our surprise, analysis of the available genomic sequences of *T. vaginalis* showed that this organism does not contain a *nox* gene encoding a typical tDBDF superfamily member, but instead contains 3 copies of a natural fusion between flavodiiron, rubredoxin and NADH:rubredoxin oxidoreductase domains (Class F FDP or FDPF according to the new classification) (**Fig. 2B**) (27). These copies have pairwise amino acid sequence identities of 75-80 % and likely arose through gene duplication, as it is common in many protozoa (34). We subsequently overexpressed and purified these three Class F FDPs from *T. vaginalis* (*Tv*FDPF1-3). After preliminary kinetic characterization of these 3 proteins, we focused our extensive biochemical and biophysical characterization on *Tv*FDPF3, as it was the most active and well-behaved. Our biochemical experiments with *Tv*FDPF3 confirmed that this natural fusion protein allows the crosstalk of all three domains, ultimately relaying electrons from NAD(P)H to O_2_. We also used single-particle electron cryo-microscopy (cryo-EM) to visualize the dimerization interface of this Class F FDP. Overall, these studies allow us to conclude that the main H_2_O-forming oxidase activity in *T. vaginalis* is catalyzed by a fusion Class F FDP we identified and characterized. Our observations suggest the activity described by Bradley and Linstead in 1988, and by Tanabe in 1979, was misassigned as a NOX-like enzyme, and in fact is a Class F FDP that catalyzes the H_2_O-forming oxidase reaction within a single polypeptide. Our spectroscopic and structural studies provide important new insight into how the Class F FDP system from *T. vaginalis* achieves extremely efficient H_2_O-forming oxidase activity using four separate redox centers.

## Results

### Identification of Class F flavodiiron proteins in *T. vaginalis* as enzyme candidates for previously reported H_2_O-forming NADH oxidase activity

Because we were interested in evolutionary innovations that allow lower organisms that lack a membrane bound ETC to maintain an optimal intracellular redox environment we explored NOXes ascribed to human protozoan parasites (4). While we easily identified previously characterized NOX in the genome of *G. lamblia* (32,33,35), the closest matches in the genome of *T. vaginalis* were three proteins, all of which were 871 amino acids in length and annotated in the GeneBank database as pyridine nucleotide-disulfide oxidoreductases or apoptosis inducing factor (TVAG_263800, TVAG_049830 and TVAG_121610) (**Supplementary Table S1**). On closer inspection, we noticed that these 3 proteins represent a natural fusion composed of an N-terminal portion homologous to a flavodiiron protein, a middle rubredoxin (Rb) domain, and a C-terminal portion that is homologous to an NADH:rubredoxin oxidoreductase (NROR) (**Fig. 2B**). Based on the recent classification all 3 fusion proteins in *T. vaginalis* are Class F FDPs (from now on we denote them as *Tv*FDPF1-3)(27) (**Fig. 2B**).

When compared to FDPAs from *T. vaginalis*, *G. lamblia* and several other organisms, the N-terminal FDP domain of *Tv*FDPF1-3 includes the previously recognized canonical sequences containing the ligands of the diiron center: Fe 1 (His82-X-Glu84-X-Asp86-His87, *Tv*FDPF3 numbering) and Fe 2 (His148-X_18_-Asp166-X_64_-His230) (**Supplementary Figure S1**). The middle domain of *Tv*FDPF1-3 is homologous to rubredoxins and the C-terminal domain of bacterial rubrerythrin (**Supplementary Figure S2**). A central feature of the middle domain of *Tv*FDPF1-3 is the presence of two pairs of Cys-X_2_-Cys sequence patterns that coordinate the iron of the Fe(SCys)_4_ center. This middle Rb-like domain is flanked at both sides by ~25-30 amino acid linkers.

NADH:rubredoxin oxidoreductases and NOXes both belong to the tDBDF superfamily of flavoenzymes allowing our initial BLAST analysis to identify *Tv*FDPF1-3 (*i.e*. the C-terminal domain of Class F FDPs [*i.e*. NROR] is homologous to NOX) (**Supplementary Figure S3**) (9). Both the GxGxxG dinucleotide-binding motif and the NAD(P)H substrate specificity loop are identifiable in the C-terminal NROR domain of *Tv*FDPF1-3 as well as in other closely related members of the tDBDF superfamily, including a “stand-alone” NROR, “stand-alone” NADH:ferredoxin oxidoreductase, and CoA-disulfide reductase (**Supplementary Figure S3**).

### Cloning and initial purification of Class F fusion flavodiiron proteins from *T. vaginalis* (*Tv*FDPF1-3)

To test whether *Tv*FDPF1-3 exhibit the H_2_O-forming oxidase activity and accept electrons directly from NADH or NADPH, the corresponding genes from *T. vaginalis* were cloned with a C-terminal Hexa-His tag into a bacterial expression vector. All 3 genes had no introns, as it is common in *T. vaginalis*, *G. lamblia* and other parasitic protozoa. When purified, all three recombinant proteins were brown in color and migrated as ~100 kDa band as judged by the SDS-PAGE analysis (**Supplementary Figure S4A-C**, **Supplementary Figure S5**). Notably, the *Tv*FDPF2 protein was prone to severe aggregation and during purification eluted in the void volume of the size-exclusion column. Based on analytical gel-filtration and assuming a globular shape, we estimated an apparent molecular weight for *Tv*FDPF1 of 302 ± 10 kDa and for *Tv*FDPF3 of 326 ± 5 kDa. The absorption spectra of all three oxidized proteins had clear features of flavin bands (350-500 nm) and a Fe(SCys)_4_ center of rubredoxin (>500 nm) at 379, 454, 475 and 568 nm for *Tv*FDPF1; 374, 451, 478 and 565 nm for *Tv*FDPF2; and 378, 451, 475 and 574 nm for *Tv*FDPF3 (**Supplementary Figure S4B-C, Supplementary Figure S5**). In the presence of excess sodium dithionite these cofactors were reduced, and the visible absorbance was almost completely bleached.

We next employed liquid chromatography–mass spectrometry (LC-MS) to determine both the molecular identity and quantity of flavin cofactors bound to *Tv*FDPF1-3. While we unambiguously identified both FAD and FMN within *Tv*FDPF1-3, the actual stoichiometries of these cofactors per protein monomer were drastically different (**Table 1**). Both *Tv*FDPF1 and *Tv*FDPF3 had full occupancy with FAD per monomer (1.02 ± 0.05 and 1.00 ± 0.12, respectively). However, FMN was present only at 0.010 ± 0.004 occupancy per monomer (~ 1 %) in *Tv*FDPF1 and 0.090 ± 0.006 (~ 9 %) in *Tv*FDPF3 (**Table 1**). *Tv*FDPF2 had poor occupancy of both FAD and FMN, which is not surprising given its poor behavior during purification. Thus far only the FMN cofactor was reported in Class A FDPs, and it seemed logical that poor occupancy of FMN in our preparations of *Tv*FDPF1-3 reflects occupancy of the N-terminal domain only. Direct addition of 300 μM FMN to the lysis buffer during purification of *Tv*FDPF1 did not improve FMN occupancy (**Table 1**).

**Table 1.**
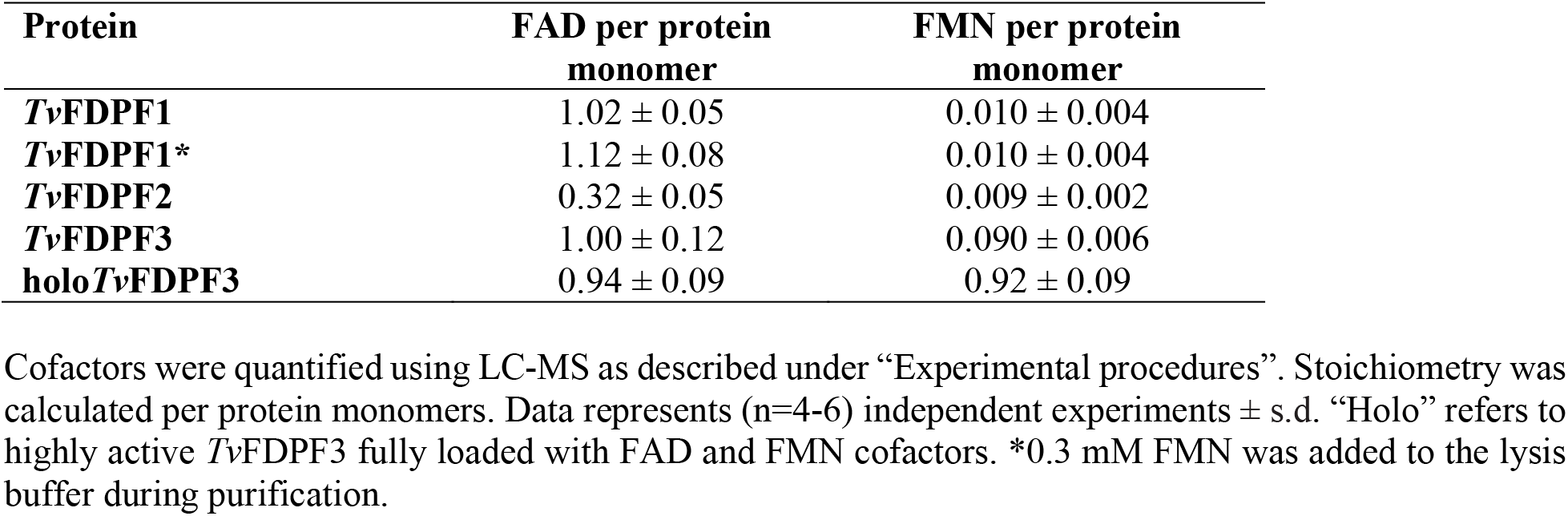
FAD and FMN quantification.

Because Fe is a part of both the diiron center and the rubredoxin-like domain, we measured iron content of all three proteins (**Supplementary Table S2**). We found that despite addition of Mohr’s salt (ammonium iron(II) sulfate) during protein expression, all purified *Tv*FDPF1-3 had substoichiometric iron occupancy (3 Fe per monomer is expected). The highest Fe occupancy of 0.82 ± 0.17 was observed for *Tv*FDPF1, while *Tv*FDPF2 and *Tv*FDPF3 had 0.41 ± 0.02 and 0.58 ± 0.10, respectively (**Supplementary Table S2**).

### Specific activity and specificity towards NADH and NADPH

Because previously reported H_2_O-forming NADH oxidase-like activities purified from *T. vaginalis* were tested with both NADH and NADPH, we tested consumption of both redox cofactors in our enzymatic assays (**Table 2**, **Supplementary Figure S4D-F**). Michaelis-Menten analysis of the reaction catalyzed by *Tv*FDPF1-3 indicates that NADH is the preferred substrate over NADPH for all three enzymes. The highest *V*_max_ of 12 ± 1 μmol min^−1^ mg^−1^ at 37 °C was observed for *Tv*FDPF3. We found that the lowest *K*_M_ for NADH among all variants was 2.9 ± 0.8 μM for *Tv*FDPF1. This value is in sharp contrast to the *K*_M_ for NADH of *Tv*FDPF3 that was 40 ± 8 μM. However, because *V*_max_ for *Tv*FDPF1 was low, the overall catalytic efficiency *k*_cat_/*K*_M_ of both *Tv*FDPF1 and *Tv*FDPF3 as well of *Tv*FDPF2 with NADH were very similar. Michaelis-Menten fitting of the NADPH data results in *K*_M_ values in the mM range, far above its physiological concentration (**Table 2**). Our results clearly support our prediction that the fusion Class F FDP from *T. vaginalis* is a “self-sufficient” enzyme that directly accepts reducing equivalents from NAD(P)H. Because *Tv*FDPF2 had low yield and was not stable, we focused further kinetic studies on *Tv*FDPF1 and *Tv*FDPF3. We also determined that *Tv*FDPF3 had little activity with NO (<2 % when compared to the NADH to O_2_ activity, **Supplementary Figure S6**).

**Table 2.** Steady-state kinetic parameters of the enzymatic reactions catalyzed by *Tv*FDPF1-3 and holo*Tv*FDPF3.

| Protein | Substrate | $K_m$<br>($\mu\text{M}$) | $V_{\max}$<br>($\mu\text{mol min}^{-1}$<br>$\text{mg}^{-1}$) | $k_{\text{cat}}$<br>($\text{s}^{-1}$) | $k_{\text{cat}}/K_m$<br>($\text{s}^{-1} \text{M}^{-1}$) |
| --- | --- | --- | --- | --- | --- |
| ***Tv*FDPF1** | NADH | $2.9 \pm 0.8$ | $4.9 \pm 1.2$ | $7.8 \pm 1.9$ | $(2.6 \pm 0.9) \times 10^6$ |
| | NADPH | $1475 \pm 486$ | $4.9 \pm 2.1$ | $7.8 \pm 3.3$ | $(5.2 \pm 2.8) \times 10^3$ |
| ***Tv*FDPF2** | NADH | $18 \pm 2$ | $6.1 \pm 0.2$ | $9.6 \pm 0.3$ | $(5.3 \pm 0.6) \times 10^5$ |
| | NADPH | $1171 \pm 208$ | $1.1 \pm 0.2$ | $1.7 \pm 0.3$ | $(1.4 \pm 0.3) \times 10^3$ |
| ***Tv*FDPF3** | NADH | $40 \pm 8$ | $12 \pm 1$ | $19 \pm 2$ | $(4.8 \pm 1.0) \times 10^5$ |
| | NADPH | $399 \pm 8$ | $4.0 \pm 0.1$ | $6.4 \pm 0.2$ | $(1.6 \pm 0.0) \times 10^4$ |
| ***holoTv*FDPF3** | NADH | $56 \pm 2$ | $291 \pm 26$ | $466 \pm 42$ | $(8.3 \pm 0.8) \times 10^6$ |
| | NADPH | $427 \pm 160$ | $11 \pm 4$ | $18 \pm 6$ | $(4.1 \pm 2.1) \times 10^4$ |

Activities were measured as described under “Experimental procedures”. Kinetic parameters represent the average of (n=4-8) independent experiments ± s.d. $k_{\text{cat}}$ values were calculated per protein dimer.

### Kinetics of oxygen consumption and H_2_O_2_ production

We next studied the reaction catalyzed by *Tv*FDPF1 and 3, by monitoring both NADH and oxygen consumption simultaneously (**Supplementary Figure S7A-B**). To our surprise there was a clear difference in NADH-to-O_2_ stoichiometry for *Tv*FDPF1 when compared to *Tv*FDPF3. With *Tv*FDPF3, two consecutive additions of 250 μM NADH were needed to completely reduce O_2_ of air-saturated buffer (~250 μM) to water. In contrast, O_2_ and NADH consumption by *Tv*FDPF1 had a 1:1 stoichiometry. The observed reaction traces support a H_2_O-forming reaction for *Tv*FDPF3 but not for *Tv*FDPF1. We tested this hypothesis by measuring H_2_O_2_ produced by either *Tv*FDPF1 or *Tv*FDPF3 with Amplex Red in a discontinuous assay (**Supplementary Table S3**). We found that *Tv*FDPF1 produces 86 ± 5 % H_2_O_2_ while *Tv*FDPF3 produces only 9.6 ± 3.7 %.

### Purification of highly active holo*Tv*FDPF3

We noticed that a minor peak but with extremely high specific activity eluted in the wash during the anion exchange purification step of *Tv*FDPF3. After modifying our purification scheme, we were able to significantly enrich and better separate this *Tv*FDPF3 fraction, which had near-stoichiometric FAD and FMN occupancy (0.94 ± 0.09 and 0.92 ± 0.09) (**Table 1**, **Fig. 3A**). The apparent molecular mass of such preparations, which we refer to as “holo*Tv*FDPF3”, was 258 ± 3 kDa, as judged by analytical gel-filtration (with a sharper appearance compared to both *Tv*FDPF1 and 3). We used BN-PAGE as an independent method to assess the molecular weight of holo*Tv*FDPF3 and observed an apparent molecular mass of 322 ± 24 kDa (**Supplementary Figure S8**).

**Figure 3.**
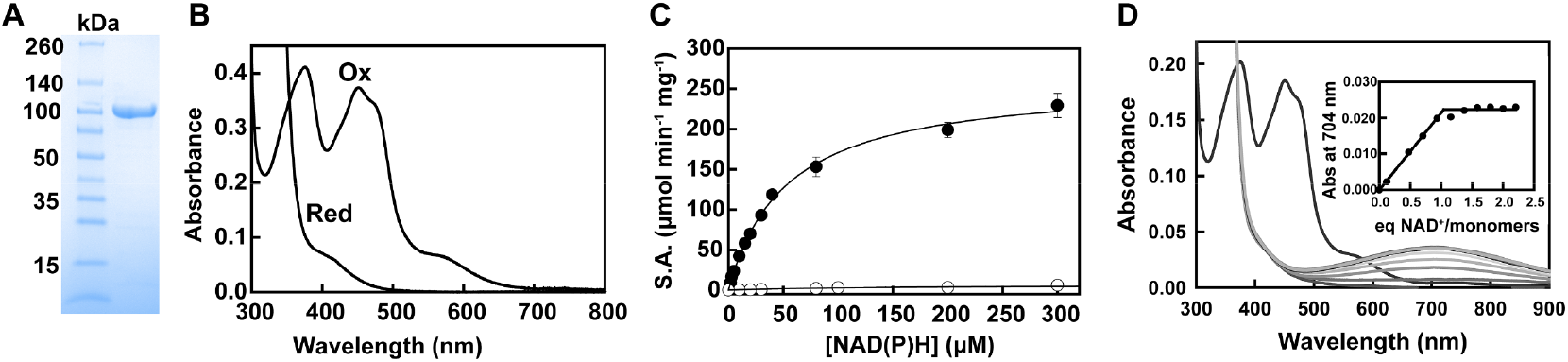
Biochemical and spectroscopic properties of highly active holoTvFDPF3. **(a)** Purified holo*Tv*FDPF3 (10 μg per lane). **(b)** UV-visible spectra of holo*Tv*FDPF3 as purified. Protein was in buffer E at 20 μM (calculated based on the molecular weight of a monomer) as purified (Ox.) and after addition of 1 mM of sodium dithionite (Red.) under aerobic conditions. **(c)** Michaelis-Menten analysis of the oxidase activity of holo*Tv*FDPF3 as described under “Experimental procedures” with NADH (filled circles) and NADPH (open circles). All kinetic parameters are summarized in Table 2. **(d)** Spectral titration of dithionite-reduced holo*Tv*FDPF3 with NAD^+^. Under anaerobic conditions holo*Tv*FDPF3 (8.18 μM of monomers) was reduced with 1 mM dithionite before different amounts of NAD^+^ were added. Spectra shown represent the starting oxidized enzyme, the dithionite-reduced enzyme and the reduced enzyme plus 0.12 - 2.2 equivalents of NAD^+^/monomer. The inset shows the absorbance change at 704 nm versus equivalents of NAD^+^/monomer.

The absorption spectrum of holo*Tv*FDPF3 had stronger features of flavin bands as well as of the rubredoxin-like center at 375, 451, 472 and 566 nm (**Fig. 3B**). Most importantly, the Michaelis-Menten analysis revealed that the V_max_ of holo*Tv*FDPF3 was 291 ± 26 μmol min^−1^ mg^−1^, a significant increase (~ 24 times) compared to the low FMN occupancy *Tv*FDPF3. The *K*_M_s for NADH and NADPH were 56 ± 2 μM and 427 ± 160 μM, respectively and *k*_cat_ (with NADH) was 466 ± 42 s^−1^ per monomer (**Table 2, Fig. 3C**). The iron content of holo*Tv*FDPF3 was 1.70 ± 0.12, which is 3 times higher compared to initially purified low activity *Tv*FDPF3 (**Supplementary Table S2**). We also used ICP-MS to characterize the metal content of holo*Tv*FDPF3 and detected a small amount of Zn^2+^ that is likely incorporated in place of Fe^2+/3+^ in the diiron site (**Supplementary Table S4**). Other metal ions were present only in negligible quantities. Finally, the H_2_O_2_ byproduct formation was two-fold lower (5.0 ± 2.7 %) than of initially purified *Tv*FDPF3 (**Supplementary Table S3**). In summary, holo*Tv*FDPF3 had more complete FAD/FMN/Fe occupancy, as reflected in its increased specific activity and we employed it in all our subsequent experiments.

### Titration of dithionite-reduced enzyme with NAD^+^

To directly demonstrate the entry point of reducing equivalents into holo*Tv*FDPF3 we performed anaerobic titration with NAD^+^ after initial reduction with dithionite (36). Oxidized NAD^+^ addition causes the appearance of a long wavelength band centered at 704 nm (**Fig. 3D**). This broad band reflects stacking of FADH_2_ and NAD^+^ planes (charge transfer complex) (36,37). This absorbance change is complete at exactly 1 equivalent of NAD^+^ per enzyme monomer (each monomer of *Tv*FDPF3 contains one FAD and one FMN). This clearly suggests that although both flavins in the protein can be chemically reduced by dithionite only one flavin binds NAD^+^ tightly. Since FMN of the Class A FDPs is not reduced by NADH, the observed behavior is consistent with the entry of NADH via FADH_2_ of the C-terminal NROR domain.

### Insights into redox centers of holo*Tv*FDPF3 by EPR spectroscopy

The architecture of the fusion FDP system presents a unique opportunity to biophysically characterize redox centers that are amenable to Electron Paramagnetic Resonance (EPR) spectroscopy. FAD and FMN absorption in the visible region overwhelm the moderate absorption of the rubredoxin center while the diiron center does not absorb in the UV-Visible range. On the other hand, oxidized and reduced flavins are diamagnetic and thus invisible by EPR. In frozen samples at low temperature signals of semiquinone radicals can be detected but are saturated at the microwave power used and do not overlap. Thus, in the “as-isolated” holo*Tv*FDPF3 protein a very strong isotropic EPR signal at *g*=4.3 could easily be detected (**Fig. 4A**). This signal is typical for the oxidized Rb-like center, of which the d^5^ ferric ion is in a high spin state (*S*=5/2). Such high spin ferric species exhibit three EPR signals from its Kramers’ doublets, of which only the | ±3/2> doublet shows an intense signal. Three almost identical *g*-values at 4.3 occur if the ratio of the spin Hamiltonian parameters E and D is above 0.3 (| E/D| >0.3). Weak signals of the two other doublets (*i.e*. only the absorption-shaped signal corresponding to the highest *g*-value) could also be detected. The intensity of the temperature corrected *g*=9.3 EPR signal of the |±1/2> doublet decreased above 4 K, indicating D = +(1.3 ± 0.6) cm^−1^ (**Fig. 4B**). In the 15 K minus 9 K difference EPR spectrum the *g*=9.8 EPR feature of the | ±5/2> doublet could be revealed. For the determination of the redox midpoint potential at room temperature holo*Tv*FDPF3 was oxidatively and reductively titrated in the presence of organic dyes by addition of potassium ferricyanide or sodium dithionite, respectively. When such samples were frozen in liquid nitrogen, the intensity of the *g*=4.3 EPR signal of the ferric state decreased upon reduction, in agreement with conversion to the EPR silent *S*=2 ferrous state of the Rb-like center (**Fig. 4C**). By fitting the intensity to the appropriate Nernst equation for a single electron process a midpoint potential of –(56±10) mV vs. H_2_/H^+^ for the Rb-like center was determined (**Fig. 5A**).

**Figure 4.**
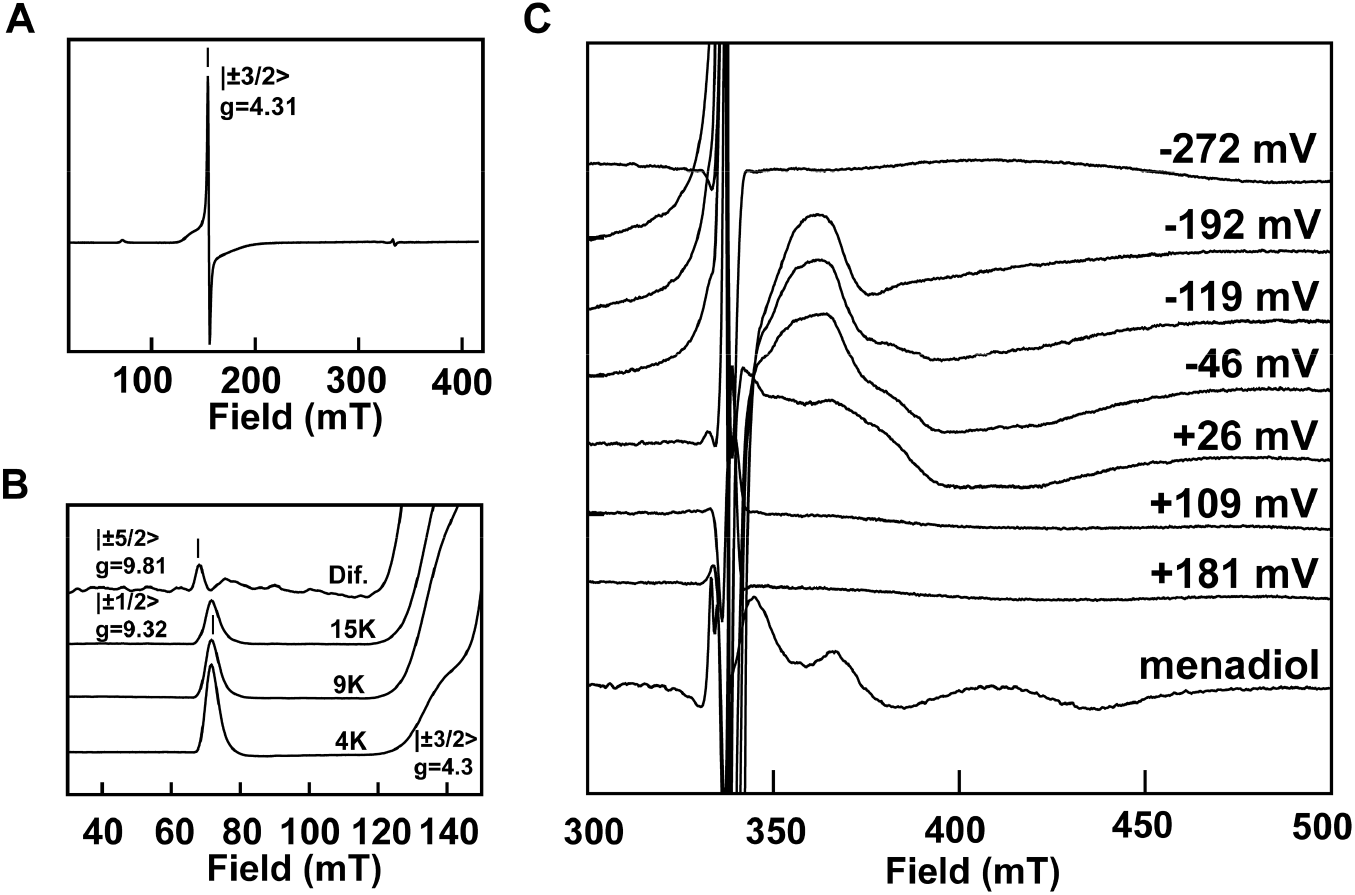
EPR analysis of holoTvFDPF3. Spectra of the ferric rubredoxin site in the as isolated enzyme at 15 K **(a)**, or at indicated temperatures (including a 15 K minus 9 K difference spectrum) **(b)**. **(c)** Spectra recorded at 7.5 K for samples poised at the indicated redox potentials (vs. H_2_/H^+^ at pH 7.5) or poised at +115 mV and treated with 50 μM menadiol (all spectra were recorded at X-band, microwave frequency 9.358 ± 0.003 GHz; modulation amplitude, 1.5 mT; 2 mW microwave power; modulation frequency 100 kHz) as described under Experimental procedures.

**Figure 5.**
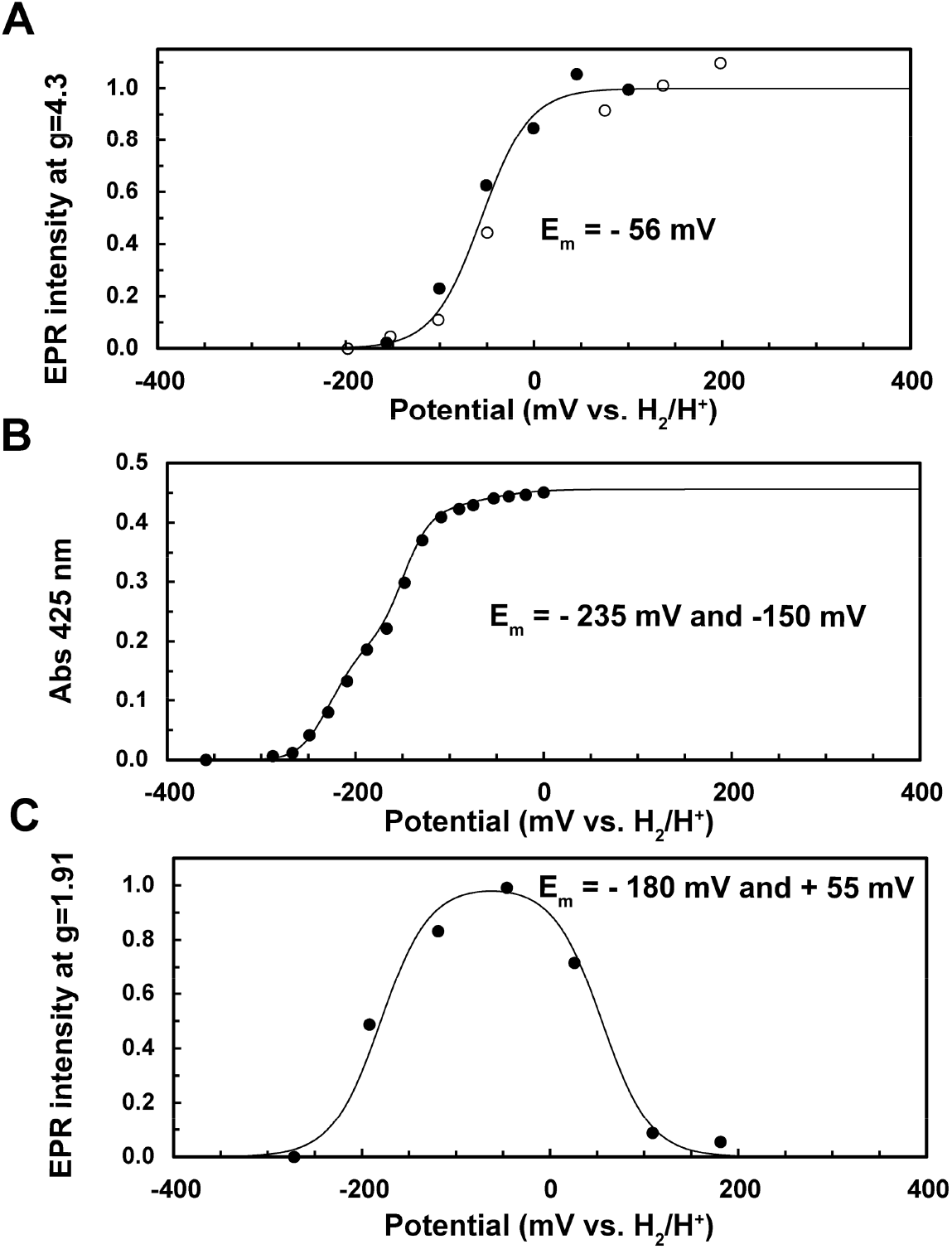
Determination of redox potentials of cofactors in holoTvFDPF3. **(a)** Intensities of the *g*=4.3 EPR signal (recorded at 77 K) of the rubredoxin site in frozen samples upon dye-mediated redox titrations (open and closed symbols refer two separate titrations). **(b)** Intensity of the mixed-valence EPR signal (amplitude at *g*=1.91) of the dinuclear iron center upon dye-mediated redox titration. EPR conditions as in Fig. 4. **(c)** Absorbance at 425 nm (corrected for contribution of the mediator cocktail in a parallel titration) reporting mainly on the presence of the oxidized flavin cofactors at room temperature. Fits to the (sum of) Nernst equation(s) are shown: rubredoxin site E_m_=-56 mV, n=1 (a), FAD, E_m_=-235 (n=1) and −235 mV (n=1), FMN, E_m_=-190 (n=1) and −110 mV (n=1), flavins each with a 0.215 absorbance plus an absorbance of 0.026 for the rubredoxin site (E_m_=-56 mV, n=1) in (c). In (b) the curve for E_m_=-180 (n=1) and +55 mV (n=1) indicates the approximate redox potentials.

The EPR detection of the diiron center and therefore the analysis of its redox chemistry is inherently more difficult. First, the diferrous and diferric states are diamagnetic or have integer spin and are not (easily) detectable. Detection of the mixed-valence dinuclear center (Fe^2+^-Fe^3+^) is hampered by the narrow temperature range at which the extremely anisotropic and large linewidth *S*=1/2 EPR signal can be detected. But at moderate microwave power at 7 K and by combining several preparations of holo*Tv*FDPF3 we could detect a rhombic signal with *g*=1.94, 1.79 and 1.53 in a sample titrated to a solution potential of+115 mV which was achieved by addition of menadiol (**Fig. 4C**). Menadiol (reduced menadione) was previously used to characterize the EPR spectrum of a Class F FDP from *Clostridium difficile* by Folgosa and colleagues (38). Due to the small quantity of the high activity *holo* enzyme and the weak EPR intensity it was not feasible to perform extensive redox titrations as in case of the Rb-like center. By combining three high activity preparations dye-mediated titrations with a total number of 7 datapoints enabled us to follow the EPR intensity of the mixed-valence EPR signal as a function of the solution potential. The diferric to mixed-valence potential was determined being +(55±30) mV (**Fig. 5B**). Line shape changes, possibly due to reduction of the nearby FMN or conformational changes, only allowed us to estimate the mixed valence to diferrous state midpoint potential to –(180±50) mV (**Fig. 5B**).

### Redox titration of holo*Tv*FDPF3 followed by the UV-Visible spectroscopy

To complete our characterization of the redox chemistry of holo*Tv*FDPF3, visible spectroscopy at room temperature was employed to estimate the midpoint potentials of FAD and FMN in the presence of 5 μM mediator concentrations in a reductive titration. Flavin visible spectra of the protein bleached due to reduction to the hydroquinone states before onset of strongly absorbing viologen signals. At a wavelength of 425 nm the contribution of the mediators was relatively low and constant. After spectral subtraction the intensity of the sum of FMN and FAD visible contributions could be followed (**Fig. 5C**). In our experiments no flavin semiquinone signals were detected. Two separate redox potential ranges with 425 nm absorbance changes were observed, which based on the nearly stoichiometric presence of FMN and FAD likely correspond to redox transitions of the individual flavins. We assigned the lower flavin potential to the FAD/FADH_2_ couple, as logical low potential entrance into the ET pathway. The best fit to the experimental data required equal potentials of −235 mV vs. H_2_H^+^ (pH 7.5) for the two n=1 transitions of FADH_2_ (**Fig. 5C**). A maximal population of 33 % semiquinone can be calculated, which in its anionic form is apparently too weak to be detected among mediator and reduced FAD and FMN signals. FMN behaved almost like an n=2 redox system with −150 mV vs. H_2_H^+^ (pH 7.5), *i.e*. extensive disproportionation of the semiquinone (max. 2 %) with estimated potentials of −110 (FMNH_2_/FMNH) and −190 mV (FMNH/FMN).

In summary, electrons within the multidomain system of *Tv*FDPF3 flow from NADH/NAD^+^ (−335 mV at pH 7.5) to FAD (−235 mV), rubredoxin (−56 mV), FMN (−150 mV) and ultimately on the dinuclear center (−180 and +55 mV, average −63 mV), the later one, in its reduced state reacts with dioxygen. We assume that the strong electrochemical driving force from NADH and FADH_2_ pushes the two times one electron transfer from the Rb-like center to the FMN within the FDP domain (see Discussion below).

### Molecular architecture of holo*Tv*FDPF3

We used single-particle cryo-EM to investigate domain organization and redox center positioning in holo*Tv*FDPF3. Two-dimensional class-averages had clear secondary structure and were approximately 80-100 Å in diameter, consistent with the expected size of a globular monomer or a portion of a higher oligomer (**Fig. 6A**). Three-dimensional reconstruction of the particle set without symmetry enforced produced a 6.6 Å resolution map (**Supplementary Figs. S9, S10**; **Supplementary Table S5**) with local resolutions ranging from 6.2 Å at the core of the particle to 7.0 Å at the periphery (**Supplementary Fig. S10A**). At this resolution, alpha helices could be clearly delineated but a model could not be constructed *de novo*. We modeled individual domains by threading the *Tv*FDPF3 sequence onto structures of homologous proteins using PHYRE2 (39). The models were then manually docked into the cryo-EM density, revealing two FDP-like domains bound in a C2-symmetric dimer interface (**Supplementary Fig. S11**). Electron density was noticeably poorer for one subunit of the dimer, potentially as a result of conformational heterogeneity produced by hinge motions at the dimer interface. We refined the density map again with C2 symmetry applied, resulting in a symmetric map with 6.8 Å nominal resolution, clear density for both subunits of the dimer, and local resolutions ranging from 5.4 Å to 6.9 Å (**Fig. 6B, Supplementary Fig. S12**). Docking homology models of the FDP-like domain into the map with C2-symmetry applied produced a model equivalent to that generated from the original map without symmetry applied. In the final C2-symmetric map, density was observed for the diiron and FMN moieties in the positions expected from previously reported structures of Class A FDPs (**Supplementary Fig. S13**).

**Figure 6.**
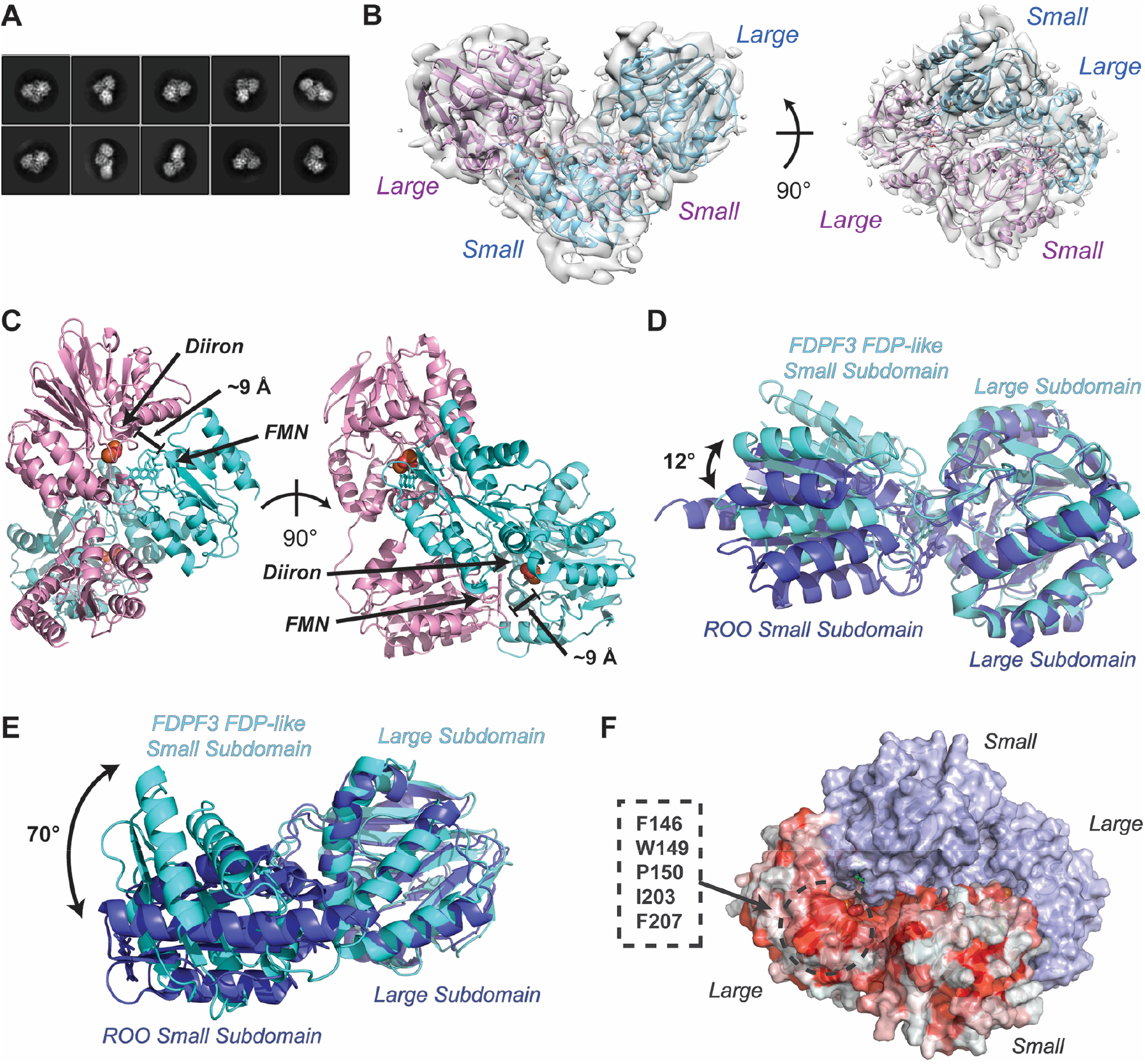
Cryo-EM structure of the FDP-like core of the full-length TvFDPF3. **(a)** Representative 2D class averaged images of holo*Tv*FDPF3 particles. **(b)** Homology models docked into the reconstructed single-particle cryo-EM C2-symmetric map. **(c)** Distance between diiron center and FMN in the holo*Tv*FDPF3 FDP-like domain dimer. **(d)** Aligned overlay of the holo*Tv*FDPF3 FDP-like domain redox center with the redox center in ROO from *D. gigas* (PDB 1E5D) (17). **(e)** Aligned overlay of one holo*Tv*FDPF3 FDP-like domain subunit with a subunit of ROO from *D. gigas*, as above. **(f)** Surface representation of the dimerized FDP-like domain model with one subunit colored by hydrophobicity. A hydrophobic patch that is concealed in the *D. gigas* ROO head-to-tail dimer is highlighted.

The FDP-like domain of *Tv*FDPF3 forms an expected “head-to-tail” dimer that positions the diiron center of one protomer 8-10 Å away from the FMN moiety bound in the neighboring subunit (**Fig. 6C**). However, the dimer interface differs from that previously reported in structures of dimeric Class A FDPs from *D. gigas*, *G. intestinalis* and *M. thermoacetica* (14,17,19). The *trans* interaction between the large subdomain of one subunit and the small subdomain of its neighbor is similar to structures reported previously (all-atom RMSD 1.570), with a 12° rotation about the redox center (**Fig. 6D**). The holo*Tv*FDPF3 dimer interface differs from that observed for Class A FDP from *D. gigas* (ROO) as a result of a 70° rotation about the linker connecting the large and small subdomains of the FDP-like core (**Fig. 6E**). This rotation substantially alters the interface between protomers (**Supplementary Fig. S14**).

## Discussion

There are substantial gaps in our understanding of core energy metabolism and redox maintenance in microaerophilic human parasites that belong to the *Excavata* supergroup (4,40). Most of their proteomes, including key metabolic enzymes, are not well characterized and are annotated primarily based on sequence homology. In this work we aimed to close this knowledge gap by biochemical and structural studies of the oxygen detoxification systems reported in *T. vaginalis* (4). The natural function of H_2_O-forming NADH oxidases (NOXes) and multicomponent flavodiiron proteins (FDPs) in microaerophilic human protozoan parasites is both regeneration of oxidized pyridine dinucleotides and constant O_2_ removal (detoxification) from the surroundings (**Fig. 1A**) (3,4,41,42). In addition, these protozoa lack enzymes typically used to combat reactive oxygen species (ROS), including catalase and superoxide dismutase (SOD), and are therefore dependent on the ability of the H_2_O-forming oxidase reaction to remove O_2_ before it can participate in the ROS-forming side reactions (3,4,42). Better understanding of the O_2_ scavenging systems are needed as these pathways were recently suggested as targets for therapeutic interventions against various human protozoan parasites (3).

The presence of a NOX in *T. vaginalis*, and its general acceptance in the literature, was based on the original work of Linstead and Bradley from 1988 (3,4,9,31). In this study we show that in *T. vaginalis* the closet homologs of a typical NOX enzyme are three very similar in amino acid sequence Class F FDPs that are currently misannotated in databases (TVAG_263800, TVAG_049830 and TVAG_121610) (**Fig. 2B**, **Supplementary Table S1**). Based on domain organization all three proteins harbor N-terminal FDP, central Rb and C-terminal NROR domains (**Fig. 2**). Of note, Smutna and colleagues previously suggested that TVAG_263800, TVAG_049830 and TVAG_121610 proteins are “self-sufficient” FDPs, however, this prediction was not tested, as only a “stand-alone” Class A FDP from *T. vaginalis* (TVAG_036010) was biochemically characterized at that time (18).

To directly test whether these genes encode “self-sufficient” FDPs, we overexpressed corresponding protein products in *E. coli* (**Supplementary Figure S4A-F).** The initial kinetic characterization of these recombinant proteins revealed that despite variable and substoichiometric loading with FMN and iron, all three enzymes are capable of using NADH or NADPH as substrates (**Tables 1–2**, **Supplementary Table S2)**. The highest V_max_ (12 ± 1 μmol min^−1^ mg^−1^) was observed for *Tv*FDPF3 with NADH. Notably, the FMN content is negatively correlated with the off-target H_2_O_2_ production as *Tv*FDPF1 (only 1% of FMN per monomer) mostly produces H_2_O_2_ (86 ± 5%) (**Supplementary Table S3**). Once the FMN loading is ~10% as in *Tv*FDPF3, the H_2_O_2_ production is significantly lower (9.6 ± 3.7%) and the NADH:O_2_ stoichiometry is 2:1. Because the FMN cofactor is a part of the FeFe-FMN diiron active site, very low levels of FMN cofactor in *Tv*FDPF1 allow only a minor fraction of the four-electron H_2_O-forming reaction to go until completion and the most of reducing equivalents are used within the C-terminal NROR domain to produce exclusively H_2_O_2_. A similar phenomenon has been reported for NROR from *Clostridium acetobutylicum*, which is a H_2_O_2_-forming NADH oxidase. When *C. acetobutylicum* NROR is mixed with fprA2 (Class A FDP) and Rb, the NROR-Rb-fprA2 system starts to catalyze an efficient H_2_O-forming oxidase reaction (23).

Next, we significantly optimized our purification scheme to obtain a highly active *Tv*FDPF3 with near-stoichiometric amounts of both FAD and FMN cofactors (holo*Tv*FDPF3) (**Table 1**, **Fig. 3A-D**). The iron content of holo*Tv*FDPF3 also improved to 1.7 ± 0.12 of Fe per monomer (**Supplementary Table S2).** Most importantly, holo*Tv*FDPF3 had increased *V_max_* of 291 ± 26 μmol min^−1^ mg^−1^ (*k*_cat_=466 ± 42 s^−1^ calculated per dimer) with *K*_M_ for NADH at 56 ± 2 μM (**Table 2**). We note that if an average of 1.3 Fe is missing from each dinuclear center, the turnover number under *V*_max_ conditions could be as high as 1310 s^−1^, or as high as 820 s^−1^, if the missing 1.3 Fe is equally distributed over both dinuclear and the rubredoxin sites (“all-or-none”). The activity we are reporting of holo*Tv*FDPF3 is similar to that reported for Class A FDP from *E. histolytica* (400 ± 30 s^−1^) (**Supplementary Table S6**). In a recent study, Class F FDP from *Clostridium difficile* was cloned and purified with iron and flavin content comparable to holo*Tv*FDPF3 and a 30-fold lower of *k*_cat_ of 16.0 s^−1^ was reported (**Supplementary Table S6**) (38).

Remarkably, the *V*_max_ we determined for holo*Tv*FDPF3 (466 ± 42 s^−1^) agrees well with the *V*_max_ number reported for the native H_2_O-forming NADH oxidase preparation from *T. vaginalis* from the study by Linstead and Bradley (470 s^−1^) (31). In the original study authors presented the purification of an H_2_O-forming NADH oxidase activity that migrated as: (*i*) two close bands of ~97-97.5 kDa on the SDS-PAGE, and (*ii*) two very similar bands during isoelectric focusing (pIs ~5.5) (31). This clearly represents a contradiction, as all known tDBDF family NOXes have molecular mass of ~50 kDa. When we compared the pIs and molecular masses of *Tv*FDPF1-3 they matched exactly the behavior of protein samples described by Linstead and Bradley (31). For example, the pI of *Tv*FDPF1 (5.25) and pIs of *Tv*FDPF2-3 (5.75 and 5.63) can explain the two very close bands during isoelectric focusing. During late stages of our study we learned that *Tv*FDPF2 was detected based on the mass spectrometric analysis in the cell lysate of *T. vaginalis* and resulted protein was in-gel stained for the NADH oxidase activity (neither specific activity nor *K*_M_ was reported) (43). In the same study, a purification of recombinant *Tv*FDPF2 for possible kinetic and spectroscopic analysis was not successful and neither *Tv*FDPF1 nor *Tv*FDPF3 were detected in cell lysates (43).

Our work complements a recent study by Folgosa and colleagues where authors determined redox potentials of redox cofactors that constitute the ET pathway within *C. difficile* FDPF (38). Here, we report a complete set of redox potentials that compose all four redox centers of the multi-domain *Tv*FDPF3 (**Fig. 5**, **Supplementary Table S7**). The redox potential of the FAD within the NROR domain (FAD, E_m_=-235 (n=1) and −235 mV (n=1)) is very similar to potentials reported for *C. difficile* FDPF and “stand-alone” NRORs from *E. coli* and *P. furiosus* (**Supplementary Table S7**). The redox potential of the Fe(SCys)_4_ center (E_m_=-56 mV, n=1) is very close to values reported in the literature for several “stand-alone” rubredoxins or the Rb-like domain of *C. difficile* FDPF (**Supplementary Table S7**). These values are very different from known redox potentials of the rubredoxin-like center of nigrerythrin and rubrerythrin (+213 to +281 mV) which is surprising as the middle domain of FDPFs is maximally homologous to rubrerythrins. The redox potential of the FMN cofactor (FMN, E_m_=-190 (n=1) and −110 mV (n=1)) is very similar to values reported for multiple Class A FDPs (**Supplementary Table S7**). Finally, in this study we were able to determine the reduction potential of the diiron center (E_m_=-180 (n=1) and +55 mV (n=1)). We note that this value is different when compared to *G. intestinalis* FDPA (E_m_= +2 and +163 mV) (**Supplementary Table S7**).

Our cryo-EM analysis revealed an unexpected architecture for the dimer interface of the N-terminal FDP core of holo*Tv*FDPF3. In contrast to previous structures of the head-to-tail “stand-alone” Class A FDPs (14,17,19), the subdomains of the FDP core exhibit a large rotation about the connecting linker, altering the dimerization interface while preserving the catalytic interface between the diiron center and FMN. We were unable to resolve electron density for Rb or NROR domains, suggesting that they may move flexibly relative to the central FDP-like core. This is not altogether unexpected, as the N-terminal FDP part is tethered to the other redox centers via an extended linker. It has been previously shown that Class A FDPs catalyze robust H_2_O formation when Rb and NROR are supplied in *trans* (12,19,20,35). The tethering of domains within *Tv*FDPF3 may increase the effective concentration of these domains and facilitate short-lived electron-shuttling intermediate conformations that were not well populated in our micrographs. Moreover, the larger-than-expected molecular masses observed in both size-exclusion chromatography and native PAGE for *Tv*FDPF1-3 suggest that the Rb-NROR domains contribute to an elongated protein shape (44). The reorganization of subdomains within the N-terminal FDP portion of *Tv*FDPF3 may hint at the interaction mechanism with the linked Rb and NROR domains. The subdomain rotation we observe in *Tv*FDPF3 exposes a hydrophobic surface that in structures of similar proteins is occluded within the dimerization interface (**Fig. 6F**). This surface is positioned approximately 35 Å away from the C-terminus of the nearest FDP-like small subdomain and approximately 10 Å away from the diiron:FMN reaction center. Although we did not observe electron density in this portion of the map, the long linker connecting the FDP-like and Rb domains could bridge this distance, permitting this surface to bind the rubredoxin domain and complete electron transfer with the NROR domain.

Both our structural insight and biochemical characterization bring up an important question about the ET pathway within the dimer of holo*Tv*FDPF3. Here, we propose that the most parsimonious flow of reducing equivalents is as follows: FAD→Fe(SCys)_4_→FMN within the subunit 1 and then the diiron center of the subunit 2 (**Fig. 7**). Spatial constraints make ET from the FAD to the Fe(SCys)_4_ center of the other subunit very unlikely. The dimerization interface holo*Tv*FDPF3 lies with the N-terminal FDP core and therefore it is unlikely that Rb and NROR from different protomers exchange electrons just like Fe-Fe and FMN centers do (**Fig. 7**). In summary, the two active sites within the dimer of “head-to-tail” FDP are formed and electrons are relayed through Rb-NROR paths separately in both subunits 1 and 2 (**Fig. 7**). Testing our hypothesis of electron transfer from NROR to the FMN and potential asymmetry in the ET pathway within active sites of the dimer of *Tv*FDPF3 is the next phase of our study.

**Figure 7.**
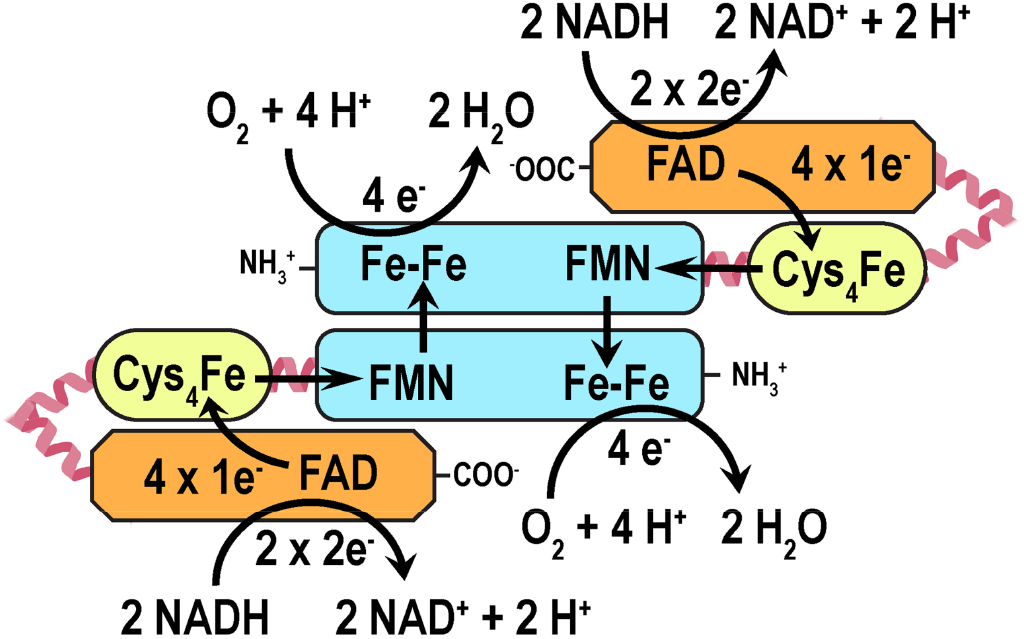
Proposed electron transport (ET) pathway within the dimer of holoTvFDPF3. Color scheme of FDP, Rb and NROR domains is the same as in Fig. 2.

### Experimental procedures

#### Materials

All chemicals were purchased from Sigma-Aldrich, VWR and Fisher Scientific unless otherwise specified. Menadiol (2-methyl-1,4-naphthalenediol) was from AK Scientific (Union City, CA).

#### Bioinformatic Analysis

The EuPathDB Bioinformatics Resource Center (https://eupathdb.org/eupathdb/) was used to browse genomic sequences of *T. vaginalis* C3 and other eukaryotic microbes (45). Multiple sequence alignments were constructed using a stand-alone version of ClustalX 2.0 and edited using BOXSHADE 3.21.

#### Cloning of TvFDPF1-3 with a C-terminal His6tag

Genomic DNA of *Trichomonas vaginalis* obtained from ATCC (Strain C-1:NIH [ATCC 30001]) was used to PCR amplify the gene sequences encoding *Tv*FDPF1-3. The following primers were used to clone TVAG_263800 (*Tv*FDPF1) contained NdeI and XhoI restriction sites (underlined): forward 5’ - TTA ATT <u>CAT ATG</u> CTT AAA ATT CAG CAA CTT ACA GAA GAT ATC - 3’ and reverse 5’ - TTA ATT <u>CTC GAG</u> AAA CAA TTC AGC AAG AAC GTG CTG TAA AGT ATG - 3’. The following primers were used to clone TVAG_049830 (*Tv*FDPF2) contained NdeI and KpnI restriction sites (underlined): forward 5’ - TTA ATT <u>CAT ATG</u> CTT AAA ATA CAG CAG CTC ACT GAA GAC - 3’ and reverse 5’ - TTA ATT <u>GGT ACC</u> TT GAA GAT TTC AGC CAT GAT GCG CTC TAA TG - 3’. The following primers were used to clone TVAG_121610 (*Tv*FDPF3) contained AseI (creates an overhang compatible with the NdeI restriction site) and XhoI restriction sites (underlined): forward 5’- TTA <u>ATT ATT</u> AAT ATG CTT AAA ATT CAG CAG CTT ACA GAA GAT - 3’ and reverse 5’ - TTA ATT <u>CTC GAG</u> AAA TAC CTC AGC AAC AAC ACG CTG CAT TGA GTG - 3’. All resulting PCR products were cloned into the pET30a expression vector (Novagen, EMD Millipore).

#### Initial expression and purification of TvFDPF1-3

All proteins in this study were overproduced in One Shot BL21 Star (DE3) chemically competent *Escherichia coli* (ThermoFisher [formerly Invitrogen], C601003). Bacterial culture was grown at 37 °C in six 2.8 L flasks, each containing 1 L of Luria-Bertani medium (Becton, Dickinson and Company, 244610) supplemented with 50 μg/ml kanamycin until the absorbance at 600 nm reached 0.4 − 0.6. At that point, the temperature was decreased to 15 °C and bacterial culture was grown for additional 2 hours. After supplementation with 400 μM ammonium iron(II) sulfate hexahydrate and 0.1 mM isopropyl β-D-1-thiogalactopyranoside bacterial cultures were grown overnight. For protein purification *E. coli* cell pellet was resuspended in 120 mL of 50 mM sodium phosphate, pH 8.0, 500 mM NaCl, 30 mM imidazole (buffer A) containing 1 mg/mL of lysozyme, four cOmplete, EDTA-free, protease inhibitor cocktail tablets (Roche), 60 μL benzonase nuclease (EMD Millipore) and 4 mM phenylmethylsulfonyl fluoride (PMSF) and were disrupted by sonication on ice for 20 min at 30-s intervals separated by 60-s cooling periods. Following centrifugation, cell lysate was passed through a 0.4 μm syringe filter, diluted to ~ 300-400 mL with buffer A and loaded onto a 20 mL HisPrep FF 16/10 column (GE Healthcare/Cytiva). All chromatographic steps were performed using an AKTA Pure (GE Healthcare) or NGC Quest 10 Plus (Bio-Rad) chromatography systems. After washing HisPrep FF 16/10 column with 500 mL of buffer A, the gradient of 30-300 mM imidazole in buffer A was applied over 300 mL. Fractions containing *Tv*FDPF1-3 had distinct brown color and were exchanged into 50 mM sodium phosphate, pH 7.5 (buffer B). In the next step, protein sample in 40 mL of buffer B was applied onto a 30 mL Source 15Q column equilibrated with 50 mM sodium phosphate, pH 7.5, 50 mM NaCl (buffer C) at a flow rate of 6 mL/min. Subsequently Source 15Q column was washed with 150 mL of buffer C and protein was eluted with 450 mL of gradient of 50-300 mM NaCl in buffer C. Size-exclusion chromatography was performed on a HiLoad 16/600 Superdex 200 (GE Healthcare/Cytiva, 28989335) or HiPrep 16/60 Sephacryl S-400 HR (GE Healthcare/Cytiva) columns equilibrated with 50 mM sodium phosphate, pH 7.5, 150 mM NaCl (buffer D) or with 50 mM HEPES-NaOH pH 7.5, 150 mM NaCl buffer (buffer E). Fractions containing protein of interest were pooled, concentrated and flash-frozen in liquid nitrogen. Only affinity purification and size-exclusion chromatography steps were used for *Tv*FDPF2 due to its low stability and tendency to precipitate.

#### Determination of molecular weight by gel-filtration

Analytical gel-filtration was performed using a Superdex 200 Increase 10/300 GL column (GE Healthcare/Cytiva) operated at 0.9 mL/min in buffer E. Calibration curve was produced using aldolase (158 kDa), conalbumin (75 kDa) and ovalbumin (44 kDa) from the high molecular weight gel-filtration calibration kit (GE Healthcare/Cytiva).

#### Blue-native polyacrylamide gel electrophoresis (BN-PAGE)

All BN-PAGE reagents were from ThermoFisher Scientific. The anode buffer was prepared using 30 mL of 20x NativePAGE running buffer and 570 mL of milliQ water. Two hundred mL of “the light” and “the dark” cathode buffers were prepared using milliQ water, 10 mL of 2X NativePAGE running buffer and 1 mL or 10 mL of cathode buffer additive. Protein samples were prepared using 4x NativePAGE sample buffer. Before the electrophoresis wells of a 3-12 % Novex Bis-Tris precast gel were filled with “the dark” cathode buffer. Samples and NativeMARK protein standards were added to assembled electrophoresis cell before “the dark” buffer was used to completely cover the inner chamber. Initial step of the electrophoresis was performed at 150 V until the dye front reached approximately one third of the gel. Then “the dark” cathode buffer in the inner chamber was replaced with “the light” cathode buffer and the run was continued till completion at 250 V. We also used recombinant H_2_O-forming NADH oxidases from *L. brevis* and *G. intestinalis* as additional molecular mass controls (11).

#### Purification of highly active holoTvFDPF3

The most active holo*Tv*FDPF3 enzyme was obtained using the protocol described above but with several modifications. *E. coli* cells were grown in 6 L of Terrific Broth (TB) medium and for the homogenization step cell pellet was resuspended in 400 mL of buffer A supplemented with 1 mg/mL of lysozyme, six cOmplete, EDTA-free protease inhibitor cocktail tablets (Roche), 120 μL benzonase nuclease and 4 mM (PMSF). Affinity chromatography was performed using a self-packed 35 mL Ni Sepharose 6 Fast Flow column. For the anion-exchanger step the Source 15Q column was equilibrated with 50 mM sodium phosphate, pH 7.5, 15 mM NaCl (buffer F). The most active holo*Tv*FDPF3 (fully loaded with both FAD and FMN and superior specific activity in the 160-210 U/mg range) was eluting as a distinct peak during the 150 mL wash step with buffer F. We note that most of the protein was eluting in the gradient as described in the section above, but that protein fraction had very low specific activity compared to the highly active holo*Tv*FDPF3. Size-exclusion chromatography was performed in buffer E.

#### UV-Visible Spectroscopy

Cary 100 and Cary 3500 UV-Visible spectrophotometers (Agilent) were used to record UV-visible spectra and perform activity assays under aerobic conditions. Anaerobic experiments were performed using Shimadzu 1900 UV-Visible spectrometer installed inside a glove box (Coy Laboratory Products). Concentration of free FAD and FMN in solution were determined spectrophotometrically using extinction coefficients for FAD at 450 nm (11.3 mM^−1^ cm^−1^) and for FMN at 446 nm (12.2 mM^−1^ cm^−1^) (46).

#### Enzymatic Assays

Enzyme activity was monitored by following the decrease of the absorbance of NADH or NADPH at 340 nm. A typical reaction mixture in 0.2 mL of the buffer E was incubated for 3 min at 37 °C before NAD(P)H (0.5-600 μM) and enzyme (0.1-10 μg) were added. An extinction coefficient (ε_340_ = 6.2 mM^−1^ cm^−1^) was used to calculate NAD(P)H oxidase activity. The *k*_cat_ values for *Tv*FDPF1-3 were calculated per dimer of the protein.

Near-simultaneous oxygen and NADH consumption were monitored using a custom-made instrument for measuring fluorescence spectra and time-resolved phosphorescence originally engineered for bioenergetics experiments with a suspension of purified mitochondria (the Mootha laboratory, Massachusetts General Hospital, Boston, MA). In the current set-up NADH was monitored by its autofluorescence (λ_ex_ = 365 nm; λ_em_ = 440 − 460 nm) and oxygen was measured using the oxygen phosphorescence sensor spot SP-PSt6-NAU (PreSens Precision Sensing GmbH, Regensburg, Germany) affixed inside a quartz cuvette. Generally, enzyme (5 − 60 μg) and NAD(P)H (100-500 μM) were added to 0.5 mL of the buffer E at 28 °C under regular oxygen tension. The instrument was operated using homemade software developed in LabView, Matlab and Arduino Software IDE. Data from each experiment was exported as a text file and analyzed using SigmaPlot 13.0.

#### Determination of FAD and FMN content

Protein samples (10-50 μM of monomers) or FAD/FMN standards (0 − 100 μM) in the buffer E or ultrapure water were incubated at 95 °C for 10 min. After centrifugation at maximum speed a 50-μL aliquot was added to 200 μL of 50%/50% methanol/acetonitrile solution and the resulting samples were subjected to the LC-MS analysis. For the LC-MS analysis, running buffer G was 5% acetonitrile, 20 mM ammonium acetate and 0.25% ammonium hydroxide, pH 9.0 while running buffer H was 100% acetonitrile. The gradient was run on a Dionex Ultimate 3000 system with an Xbridge amide column (2.1 x 100 mm, 2.5 μm particle size) at 220 μL/min flow rate, and started at 85% H for 0.5 minutes, ramped to 35% H over the next 3.5 minutes, ramped to 2% H over the next 2 minutes, held at 2% H for 1 minute, ramped to 85 % H for 1.5 minutes, then held at 85% H for 1.5 minutes (and ramping to 420 μL/min). The total run time was 12 minutes. The MS analysis was performed on a Thermo QExactive Orbitrap mass spectrometer operated in polarity switching mode with a scan range of 70 − 1000 *m/z* and a resolving power of 70,000 at 200 *m/z*. The resulting data for FAD and FMN were processed using Thermo Xcalibur software.

The FAD/FMN content was also analyzed on an Agilent 6495 QqQ with jet stream source coupled to an Agilent 1290 LC stack with an Agilent HILIC-Z (2.1 x 150 mm) column at the Scripps Center for Metabolomics (Department of Chemistry, TSRI). The mobile phase was composed of buffer I = 10 mM ammonium acetate, 5 μM medronic acid, pH = 9 and buffer K = 90:10 acetonitrile/water, 10 mM ammonium acetate, 5 μM medronic acid, pH = 9. The gradient started at 98% K (0 − 1 min) decreasing to 40% K (1 − 5 min) and was followed by an isocratic step (5 min – 7 min) before a 5 min post-run for column re-equilibration. The flowrate was set to 250 μL/min and the sample injection volume was 5 μL. Operating in negative ion mode, the source conditions were as follows: drying gas temperature set to 200 °C with a flowrate of 11 L/min, the sheath gas temp was 300 °C with a sheath gas flowrate of 12 L/min, the neb pressure set to 35 psi, cap voltage set to 2500V and nozzle voltage set to 1500V. Data was processed using Agilent MassHunter Quantitative analysis software. All samples had values that were within the range of the calibration standards.

#### Iron determination with Ferene

Proteins were diluted in buffer E to 10-40 μM of monomers. Iron standards were prepared using Mohr’s Salt (ammonium iron(II) sulfate hexahydrate) in ultrapure water. Subsequently 100 μL of 1% hydrochloric acid were added to 100-μL aliquots of protein samples or iron standards and tubes were incubated at 80 °C for 10 min. Next, 500 μL of 7.5% ammonium acetate were added with subsequent addition of 100 μL of 4% ascorbic acid, 100 μL of 2.5% sodium dodecylsulphate and 100 μL of iron chelator 3-(2-pyridyl)-5,6-bis(5-sulfo-2-furyl)-1,2,4-triazine disodium salt (samples were vortexed after each addition). After centrifugation at 13000x*g*, 800 μL of each sample were transferred to 1 mL plastic cuvettes to record absorbance at 592 nm.

#### Determination of H_2_O_2_ production

H_2_O_2_ production was monitored in a discontinuous assay. Large excess of the protein (5-30 μg) was added to the assay mixture that contained 110 μM NADH in 0.3 mL of buffer E and allowed to run at room temperature for 5 min (to establish full conversion of NADH to H_2_O_2_/H_2_O). Aliquots of 50 μL were taken and added to another 50 μL of Buffer E supplemented with 2 μL of HRP (Abcam, ab102500), and 2 μL Amplex Red (Abcam, ab102500). In parallel a calibration curve with known amounts of H_2_O_2_ standards was constructed. Ten minutes later after incubating the assay mixture in a clear 96-well plate, the absorbance at 600 nm was recorded using EnVision 2103 plate reader.

#### Metal content determination by ICP-MS

The ICP-MS analysis was done on a Thermo Scientific iCAP RQ ICP-MS in the Environmental and Complex Analysis Laboratory (UCSD). Protein samples were prepared at 0.5 μM of active sites (10 mL) in 2 % Trace Metal Nitric Acid and analyzed directly. The analysis was conducted in Kinetic Energy Discrimination (KED) mode monitoring 45Sc and 89Y as internal standards.

#### EPR spectroscopy

The midpoint potentials of the rubredoxin and dinuclear iron centers of holo*Tv*FDPF3 were determined from EPR signal intensities of the oxidized (as purified) and mixed-valence state, respectively. About 5 mg/mL holo*Tv*FDPF3 in (end volume 2 mL for the reductive titration with sodium dithionite and 1.5 mL for the oxidative titration with potassium ferricyanide) 50 mM HEPES-NaOH, pH 7.5, were stirred under anaerobic conditions at 298 K. The solution potential was measured with an InLab ARGENTHAL (Mettler, Germany) microelectrode (Ag/AgCl, +207 mV vs. H_2_/H^+^ with in-built Pt counter electrode) in the presence of phenazine ethosulfate, methylene blue, resorufin, indigo carmine, 2-hydroxy-1,4-naphthoquinone, *N*,*N*,*N*,*N*-tetramethyl-p-phenylendiamine, sodium anthraquinone-2-sulfonate, phenosafranin, safranin T, neutral red, benzyl- and methylviologen (all at final concentration of 20 μM). After adjustment of the potential by microliter additions of the sodium dithionite or potassium ferricyanide and 5 min equilibration, samples were withdrawn, removed from the anaerobic glove box in EPR tubes closed with ID 3 mm x OD 7 mm natural rubber tubing with 5 mm OD acrylic glass round stick. Samples were stored in liquid nitrogen until EPR spectra were recorded. For the menadiol-treated sample 300 μL of 7 mg/mL holo*Tv*FDP3 with the first five mediators (at 10 μM) were titrated to a potential of +115 mV vs. H_2_/H^+^ with sodium dithionite. Thereafter menadiol was added to a final concentration of 50 μM. EPR spectra were recorded with a digitally upgraded Bruker Elexsys E580 X band spectrometer with a 4122HQE cavity, an Oxford Instruments ESR 900 helium flow cryostat and the cryocooling system composed of a Stinger (Cold Edge Technologies) closed-cycle cryostat linked to an F-70 Sumitomo helium compressor. EPR tubes were produced by a local glassblower from Ilmasil PN tubing with OD 4.7 mm and 0.5 mm wall thickness obtained from Qsil (Langewiesen, Germany).

#### UV-visible redox titrations

*Tv*FDPF3 (1 mg in 1 mL of 50 mM HEPES-NaOH, pH 7.5) was completely reduced by addition of sodium dithionite (1 mM final concentration). Increments of a 0.2 μM NAD^+^ in the same buffer were added. The UV-visible redox titration using a mixture of redox mediators (see section above) was performed by monitoring the absorbance at 425 nm using a Shimadzu 1900 UV-Visible spectrometer installed inside the Coy glove box.

#### Half reaction of TvFDPF3 in presence of NONOate

A stock solution (10 mM) of diethylamine NONOate sodium salt hydrate (Sigma-Aldrich, Taufkirchen, Germany) was prepared by dissolving 3.9 mg in 245 μL of deoxygenated KOH (prepared by bubbling nitrogen in the anaerobic tent). Transfer of electrons from NADH to NO in the reaction catalysed by *Tv*FDPF3 was monitored at 340 nm using a Shimadzu UV-310 spectrophotometer in the Coy tent. To release NO the assay buffer, NADH (200 μM) and NONOate (100 μM) were mixed and incubated for 5 min. Next, 120 μg of the enzyme were added, and absorbance or spectral changes were followed as a function of time. Samples where enzyme or NONOate were omitted served as controls.

#### Cryo-EM sample preparation and data collection

Purified highly active holo*Tv*FDPF3 was diluted to 0.8 mg/mL using buffer E and 4 μL was applied to a glow-discharged Quantifoil 200 mesh 1.2/1.3 Cu holey carbon grid (Electron Microscopy Sciences, Hatfield, PA). Excess sample was removed by blotting with a Vitrobot Mark IV (ThermoFisher) for 5 s at +15 blotting force in an environment held at 22 °C and 100% relative humidity, then the grid was frozen by plunging into liquid ethane. The frozen grid was imaged at 300 kV with a Titan Krios microscope (ThermoFisher) with a slit energy filter (Gatan) set to 20 eV. Images were acquired in counting mode (105,000x nominal magnification, 0.825 Å pixel size) on a K3 direct electron detector (Gatan). Acquisitions were stored as 51-frame dose-weighted movies with defocus ranging from −1.3 to −2.5 μm. Each movie was collected with a 1.8 s total exposure time, a 30.3 e^−^ Å^−2^ frame^−1^ specific dose, and a 54.5 e^−^ Å^−2^ overall electron dose. Serial-EM 3.8.6 was used to automate multi-shot image acquisition, and 7,398 total movies were collected (47).

#### Cryo-EM data processing

The holo*Tv*FDPF3 image dataset was processed using CryoSPARC 3.2.0 to pick an initial particle set and RELION 3.1.3 to curate particles and refine maps (**Supplemental Fig. S9**) (48,49). All micrographs were imported into CryoSPARC and preprocessed using the CryoSPARC internal Patch Motion Correction and Patch CTF Estimation jobs. An initial particle set was generated by blob picking particles between 80 Å and 180 Å in diameter from all micrographs. These particles were twice sequentially subjected to 2D classification, and the best classes selected. The micrographs were then manually curated to remove images with minimum CTF fits of 9 Å or greater. After curation, 6,938 of the original 7,398 movies remained under consideration. The 2D classes selected from blob picking were used to train a Topaz model, and the model was used to pick particles from the curated micrographs (50). Duplicate particles were removed by excluding particles with a center within 125 Å of another particle. The coordinates of the remaining 1,511,632 particles were then encoded into star files for import into RELION using a custom python script, made available at https://github.com/tribell4310/reliosparc. The original set of 7,398 movies was separately imported into RELION and preprocessed using RELION’s implementations of MotionCor2 1.2.6 and CTFFIND 4.1.13 (51,52). STAR files containing the coordinates of particles selected in cryoSPARC were then imported into RELION and used to extract particles from the RELION-preprocessed movies with a 256 pixel box size and no binning. The particle stack was subjected to three sequential rounds of 2D classification, with the best classes selected in each iteration. The particles were re-centered by 3D auto-refinement using an initial model generated *de novo* in RELION from a larger set of curated particles. The re-centered particles were then re-extracted from micrographs with a 256 pixel box size and no binning using the refined particle coordinates. These particles were subjected to an additional cycle of 2D classification to remove any particles that would not align well after re-centering. The best classes were then 3D auto-refined, and the resulting volume was used to create a mask with a 9 pixel extension and a 7 pixel soft edge. The mask was then applied during 3D classification of the particle set into 6 classes using a *tau_fudge* parameter of 20, as has previously been used to resolve structural features in small targets (53). Two density types emerged from 3D classification: a compact map with well-resolved features and a low-resolution extended map with poor density, which may correspond to a more flexible conformation of the protein. Subsequent masked 3D classification of the compact classes did not improve the quality of the map, so all 101,628 particles in the compact conformation were included in the final particle stack. The compact conformation particles were auto-refined twice with progressively tighter masks, first with a mask with a 9 pixel extension and a 7 pixel soft edge, and then with a 7 pixel extension and a 7 pixel soft edge. The last masked refinement was then subjected to three sequential rounds of RELION’s CTF refinement protocol, producing a map with 7.0 Å overall resolution. Finally, the map was sharpened by applying a B-factor of −534.7, estimated during the RELION postprocessing protocol. The final sharpened map had an overall resolution of 6.6 Å. After modeling indicated likely C2 symmetry in the map, the set of 130,196 particles from 2D classes represented in the C1 map was three-dimensionally classified with C2 symmetry applied, a 150 Å diameter spherical mask, a *tau_fudge* factor of 20, and the C1 refined map lowpass filtered to 50 Å as an input template. Two similar compact classes emerged and were subjected to a subsequent round of unmasked 3D classification. A homogeneous class containing 53,709 particles was identified and 3D auto-refined with a mask (6 pixel extension, 4 pixel soft edge), then CTF refined using the same protocol as above, resulting in a 6.6 Å unsharpened map. The map was sharpened with a B-factor of −658.7 using the RELION postprocessing protocol, producing a final map with 6.8 Å nominal resolution. Local resolutions were estimated using RELION’s integrated local resolution estimation algorithm (54).

#### Model Building

Because the resolution of the final maps was insufficient to build atomic models *de novo*, the model was constructed using homology models from individual domains. Homology models were generated using the PHYRE2 server, and with 92% of the *Tv*FDPF3 sequence covered between three models (39). Residues 2-366 (N-terminal FDP-like domain) were modeled onto the PDB 1VME structure (Class A FDP from *Thermotoga maritima*), residues 428-472 (rubredoxin domain) were modeled onto the PDB 1LKO structure (rubrerythrin from *Desulfovibrio vulgaris*), and residues 482-871 (the C-terminal NADH:rubredoxin oxidoreductase domain) were modeled onto the PDB 3NTA structure (NADH-dependent persulfide reductase from *Shewanella loihica*). Appropriate ligands were included in the homology models based on their conserved binding pockets in homologous structures. The map was consistent with a dimer of FDP-like domains, and models were built by manually placing homology models in density using UCSF Chimera (55), with the N- and C-terminal subdomains of the FDP-like domain separated to facilitate the large rotation between the subdomains. Once the subdomains were confidently placed, we verified that the domain placements were topologically reasonable by modeling the connecting linker region using the nextgeneration kinematic closure loop modeling protocol in Rosetta (56). The model was refined to the C2-symmetric map using a single application of real-space refinement in Phenix (57). Structural alignments comparing *Tv*FDPF3 with the head-to-tail dimer form of rubredoxin:oxygen oxidoreductase from *D. gigas* (PDB 1E5D) were performed using PyMol (58). Figures were generated using UCSF Chimera and PyMol (55,58).

## Data availability

The Cryo-EM maps of *Tv*FDPF3 have been deposited into the EMDB (accession #: EMD-25790, EMD-25787), and the underlying particle images have been deposited with EMPIAR (accession #: EMPIAR-10895). The docked homology models have been deposited in Zenodo (DOI: 10.5281/zenodo.5795907). All the other data are contained within this manuscript.

## Supporting information

This article contains supporting information.

## Acknowledgements

Cryo-EM data were collected at the Harvard Cryo-EM Center for Structural Biology at Harvard Medical School. We thank Xue Fei (Massachusetts Institute of Technology) and Michelle Fry (MGH) for helpful feedback during cryo-EM map refinement.

## Author contributions

V.C. and V.K.M. conceived the study. V.C. cloned initial DNA constructs. E.N.A. with the help from V.C. and M.L.H. performed protein purification and protein characterization experiments. B.R. and A.J.P. designed and performed EPR and redox titrations experiments. B.R. performed activity assays with nitric oxide. T.A.B. and L.H.C. determined the cryo-EM structure. O.S.S. contributed to the mass spectrometry analysis. M.A.Y. assisted in oxygen consumption measurements. V.C. and E.N.A. analyzed the data and wrote the manuscript with the exception of the EPR spectroscopy section (co-written by B.R., A.J.P.) and the structural biology section (co-written by T.A.B., L.H.C.). All authors contributed to editing the manuscript and approved the final version. V.K.M. supervised initial experiments of this study. V.C. supervised all aspects of the study.

## Funding and additional information

This work was supported by grants from the National Institutes of Health (R00GM121856 and R35GM142495 to V.C., R35GM142553 to L.H.C., F32GM133047 to O.S.S., R35GM122455 and T-R01GM099683 to V.K.M., R00AG042026 and R01AA027097 to M.A.Y.) and the Helen Hay Whitney Foundation (to T.A.B.). V.K.M. is an Investigator of the Howard Hughes Medical Institute. A.J.P. acknowledges the DFG and the government of Rhineland-Palatinate for the upgrade of the EPR spectrometer.

## Abbreviations and nomenclature

ETC: electron transport chain
ET: electron transport
FDP: flavodiiron protein
NROR: NADH:rubredoxin oxidoreductase
Rb: rubredoxin

## Unpublished observations and personal communications

None

