## Supplementary Materials for "A natural fusion of flavodiiron, rubredoxin, and NADH:rubredoxin oxidoreductase domains is the highly efficient water-forming oxidase of *T. vaginalis*"

[1] Scintillon Institute, San Diego, CA 92121, USA
[2] Department of Molecular Biology, Massachusetts General Hospital, Harvard Medical School, Boston, MA 02142, USA
[3] Faculty of Chemistry, University of Kaiserslautern, Kaiserslautern, D-67663, Germany
[4] Howard Hughes Medical Institute, Massachusetts General Hospital, Boston, MA 02142, USA
[5] Department of Bioengineering, Northeastern University, Boston, MA 02115, USA
[6] Department of Chemistry, The Scripps Research Institute, La Jolla, CA 92037, USA
[Δ] Present position: CW Associate Scientist, Amgen, Thousand Oaks, CA, 91320

[*]corresponding author: Valentin Cracan

**This PDF file includes:**

**Supplementary Tables S1-S7.**

**Supplementary Figures S1-S14.**

**Supplementary References.**

### SUPPORTING TABLES

**Supplementary Table S1.**

Flavodiiron proteins found in human microaerophilic parasite *T. vaginalis.*

| Accession ID | Current annotation in databases | FDP Class | Name used in this study | Mol. mass (monomer)*, kDa | pI* |
|---|---|---|---|---|---|
| TVAG_263800 | disulfide oxidoreductase, putative | Class F | *Tv*FDPF1 | 95.5 | 5.25 |
| TVAG_049830 | disulfide oxidoreductase, putative | Class F | *Tv*FDPF2 | 94.9 | 5.75 |
| TVAG_121610 | apoptosis inducing factor, putative | Class F | *Tv*FDPF3 | 96.0 | 5.63 |
| TVAG_036010 | A-type flavoprotein | Class A | *Tv*FDPA | 46.3 | 7.95 |

*both values were computed using Expasy Swiss Bioinformatics Resource Portal. Class F FDPs from *T. vaginalis* (*Tv*FDPF1-3) are currently misannotated in the databases. Molecular masses, pIs, and the $V_{max}$ of the most active holo*Tv*FDPF3 (see Table 2) from this study are strikingly similar to biochemical properties of the native *T. vaginalis* $H_2O$-forming NADH oxidase activity purified by Linstead and Bradley in 1988 (1). M.M.- molecular mass; pI – isoelectric point.

**Supplementary Table S2.**

Iron content quantification.

| Protein | Fe per protein monomer | % loading (100% = 3 Fe per monomer) |
|---|---|---|
| ***Tv*FDPF1** | 0.82 ± 0.17 | 27 ± 6 |
| ***Tv*FDPF2** | 0.41 ± 0.02 | 14 ± 1 |
| ***Tv*FDPF3** | 0.58 ± 0.10 | 19 ± 3 |
| **holo*Tv*FDPF3** | 1.70 ± 0.12 | 57 ± 4 |

Iron content was quantified as described under "Experimental procedures". Stoichiometry was calculated per protein monomers. Data represents (n=4-6) independent experiments ± s.d.

**Supplementary Table S3.**

$H_2O_2$ production by *Tv*FDPF1-3.

| **Protein** | **$H_2O_2$ production, (%)** |
|---|---|
| ***Tv*FDPF1** | 86 ± 5 |
| ***Tv*FDPF2** | 25 ± 2 |
| ***Tv*FDPF3** | 9.6 ± 3.7 |
| ***holoTv*FDPF3** | 5.0 ± 2.7 |
| ***Lb*NOX** | 0.6 ± 0.1 |

$H_2O_2$-formation was monitored in AmplexRed-based assay at 37ºC as described under "Experimental procedures". Estimated $H_2O_2$ formation is shown as % of the reaction of NADH to $NAD^+$ conversion based on (n = 5) independent experiments ± s.d.

**Supplementary Table S4.**

Metal content of holo*Tv*FDPF3 determined by ICP-MS.

| Metal | Metal per protein monomers |
|---|---|
| **Fe** | 2.24 ± 0.03 |
| **Cr** | 0.001 ± 0.00004 |
| **Mn** | 0.0091 ± 0.00015 |
| **Cu** | -0.00104 ± 0.00009 |
| **Co** | 0.00081 ± 0.00003 |
| **Zn** | 0.110 ± 0.001 |

Metal content was determined in by Inductively Coupled Plasma Mass Spectrometry (ICP-MS) in protein samples diluted to 0.5 μM of protein monomers.

**Supplementary Table S5.**

Cryo-EM data collection and processing statistics.

| | | |
|---|---|---|
| **Accessions** | | |
| | **C1 Symmetric Map** | **C2 Symmetric Map** |
| **EMDB (EM Maps) ID** | EMD-25790 | EMD-25787 |
| **EMPIAR (EM Images) ID** | EMPIAR-10895 | |
| **Zenodo (Docked Homology Models) Accession** | 10.5281/zenodo.5795907 | |
| **Data Collection and Processing** | | |
| **Microscope** | Titan Krios | |
| **Camera** | K3 | |
| **Magnification** | 105,000 x | |
| **Voltage (kV)** | 300 | |
| **Total Electron Dose ($e^- / Å^2$)** | 54.5 | |
| **Defocus range (µm)** | -1.3 to -2.5 | |
| **Pixel size (Å)** | 0.825 | |
| **Micrographs collected** | 7,398 | |
| **Final particles** | 101,628 | 53,709 |
| **Symmetry** | C1 | C2 |
| **Resolution (Å, FSC 0.143)** | 6.6 | 6.8 |

**Supplementary Table S6.**

Activities of representative flavodiiron proteins.

| Protein | oxidase activity*, $s^{-1}$ | NOase activity, $s^{-1}$ |
|---|---|---|
| **Class A** *G. intestinalis* (2) | 37.7 ± 8.3 | 0.2 |
| **Class A** *E. histolytica* (3) | 400 ± 30 | 1.7 ± 0.4 |
| **Class A** *T. maritima* (4) | 4 | 0.05 |
| **Class A** *M. thermoacetica (5)* | 50 | 48 |
| **Class A** *D. vulgaris (5)* | 24 | 19 |
| **Class A** *C. acetobutylicum (6)* | 5 | 34 |
| **Class F** *C. difficile* (7) | 16.0 ± 1.3 | 0.20 ± 0.01 |
| **Class F** *T. vaginalis holoTv*FDP3 (this study) | 466 ± 42 | N.D. |

*Measured as $O_2$ or NADH consumption. In most studies $k_{cat}$ are reported for an arbitrary substrate concentration and not necessary under $V_{max}$ conditions. In all assays for recombinant Class A FDPs non-physiological Rb/NROR protein partners were used (NADH:flavorubredoxin oxidoreductase and truncated rubredoxin domain of *E. coli* flavorubredoxin). N.D.- not detected.

**Supplementary Table S7.**

Redox potentials of representative rubredoxins, rubrerythrins, NRORs and FDPs*.

| Protein | E1 FAD (FADox-FADsq) | E2 FAD (FADsq-FADred) | E1-FMN (FMNox-FMNsq) | error | E2-FMN (FMNsq-FMNred) | error | Rb Fe3+ to Fe2+ | E1-FeFe 2Fe3+ to Fe3+/2+ | error | E2-FeFe Fe3+/2+ to 2Fe2+ | error | Ref. |
|---|---|---|---|---|---|---|---|---|---|---|---|---|
| Rubredoxin *C. pasteurianum* | | | | | | | -57 | | | | | (8) |
| Rubredoxin *B. methylotrophicum* | | | | | | | -40 | | | | | (9) |
| Rubredoxin *C. tepidum* | | | | | | | -87 | | | | | (10) |
| Rubredoxin *H. mobilis* | | | | | | | -46 | | | | | (11) |
| Rubredoxin *P. furiosus* | | | | | | | 0 | | | | | (12) |
| Rubredoxin *D. vulgaris* | | | | | | | 0 | | | | | (13) |
| Rubredoxin *D. gigas* | | | | | | | 6 | | | | | (14) |
| Rubredoxin Domain of FlRd (Class D FDP) *E. coli* | | | | | | | -123 | | 15 | | | (15) |
| Rubrerythrin *D. vulgaris* | | | | | | | 230 | | | | | (13) |
| Rubrerythrin *D. vulgaris* | | | | | | | 281 | 339 | | 246 | | (16) |
| Nigrerythrin *D. vulgaris* | | | | | | | 213 | 300 | | 209 | | (16) |
| NROR *P. furiosus* | -173 | | | | | | | | | | | (17) |
| NROR *E. coli* (FlRd-Red) | -220 | -260 | | 15 | | 15 | | | | | | (15) |
| FlRd (Class B FDP) *E. coli* | | | -40 | 15 | -130 | 15 | -123 | | 15 | | | (15) |
| FDP part of FlRd (no Rb domain) *E. coli* | | | 0 | | -150 | | | | | | | (15) |
| FprA (Class A FDP) *M. thermoacetica* | | | -117 | 10 | -220 | 10 | | | | | | (18) |
| Hrb (Class B FDP) *M. thermoacetica* | | | -121 | 10 | -121 | 10 | -30 | | 10 | | | (18) |
| Class A FDP *G. intestinalis* | | | -66 | 15 | -83 | 15 | | 163 | 20 | 2 | 20 | (19) |
| Class A FDP *T. vaginalis* | | | 25 | | 25 | | | | | | | (20) |
| Class A FDP *E. histolytica* | | | -55 | | -140 | | | | | | | (3) |
| Class F FDP *C. difficile* | -250 | -250 | -170 | | -170 | | -110 | | | | | (7) |
| Class F FDP *T. vaginalis* | -235 | -235 | -190 | | -110 | | -56 | 55 | 30 | -180 | 50 | This study |

*Not for all studies error values were available.

### SUPPORTING FIGURES

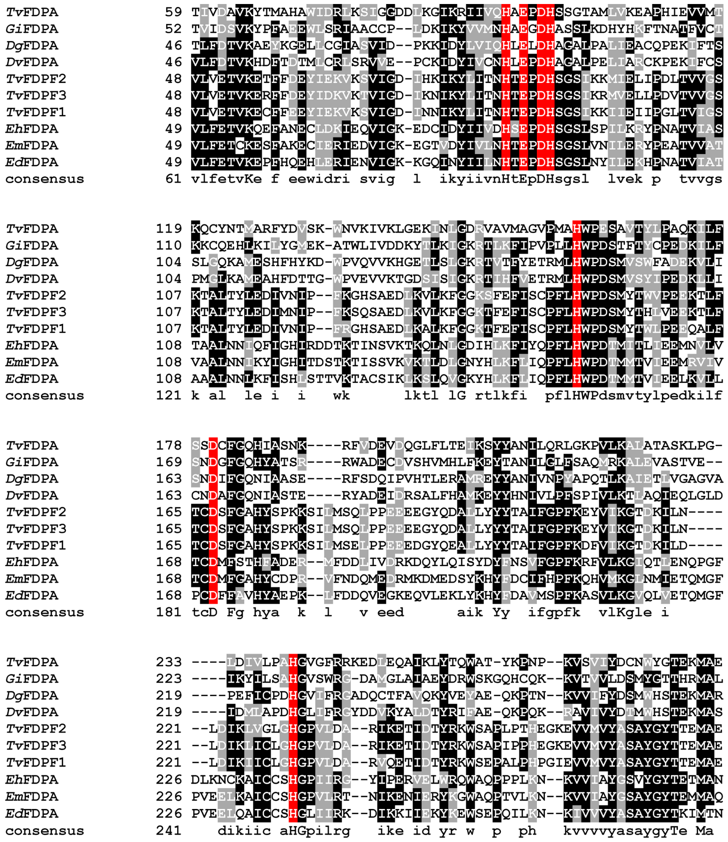


***Supplementary Figure S1.*** Multiple sequence alignment of the N-terminal sequences of Class F flavodiiron proteins from *T. vaginalis* (TvFDPF1-3) (TVAG_263800, TVAG_049830, TVAG_121610) and "stand-alone" Class A FDPs from *T. vaginalis* (TVAG_036010) and *Giradia intestinalis* (XP_001707670), *Entamoeba histolytica* (XP_651627), *Entamoeba dispar* (XP_001738262), *Entamoeba moshkovskii* (CAI11385), *Desulfovibrio gigas* (WP_021760300) and *Desulfovibrio vulgaris* (WP_010940443). Conserved residues which are important for the binding of both irons of the diiron center are highlighted in red: Fe 1 (His82-X-Glu84-X-Asp86-His87); Fe 2 (His148-$X_{18}$-Asp166-$X_{64}$-His230.

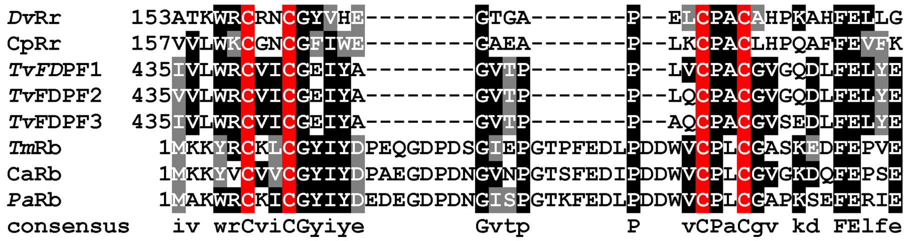


***Supplementary Figure S2.*** Multiple sequence alignment of the rubredoxin-like domain of *Tv*FDPF1-3, "stand-alone" rubredoxins from *Thermotoga maritima* (AAD35743), *Clostridium acetobutylicum* (Q9AL94) and *Pyrococcus abyssi* (WP_010868015) and the C-terminal domains of rubrerythrin from *Desulfovibrio vulgaris* (WP_010940353) and *Clostridium perfringens* (WP_164819022). Conserved cysteine residues which form the $Fe(SCys)_4$ center are highlighted. In both *Tv*FDPF1-3 and rubrerythrin the conserved sequence is CXXC-$X_{12}$-CXXC, which is different from the one found in a classical rubredoxin CXXC-$X_{29}$-CXXC.

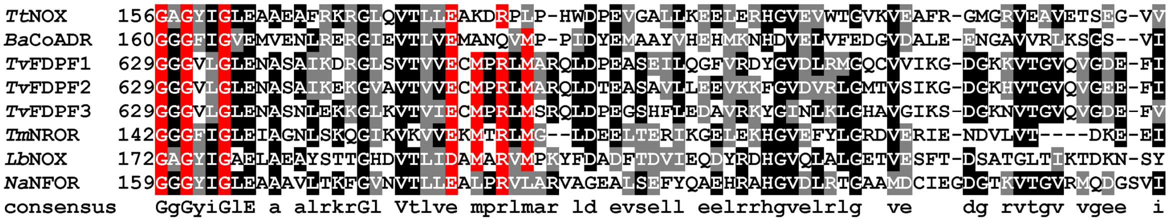


***Supplementary Figure S3.*** Multiple sequence alignment of the C-terminal NADH:rubredoxin oxidoreductase-like domains of *Tv*FDPF1-3, "stand-alone" NADH:rubredoxin oxidoreductase from *Thermotoga maritima* (WP_004080954); NADH:ferredoxin oxidoreductase from *Novosphingobium aromaticivorans* (ABD24664); $H_2O$-forming oxidases from *Lactobacillus brevis* (BAN07126) and *Thermus thermophilus* (WP_011173859) and CoA-disulfide reductase from *Bacillus anthracis* (WP_000087591). Both the dinucleotide-binding motif and NAD(P)H substrate binding loop are highlighted.

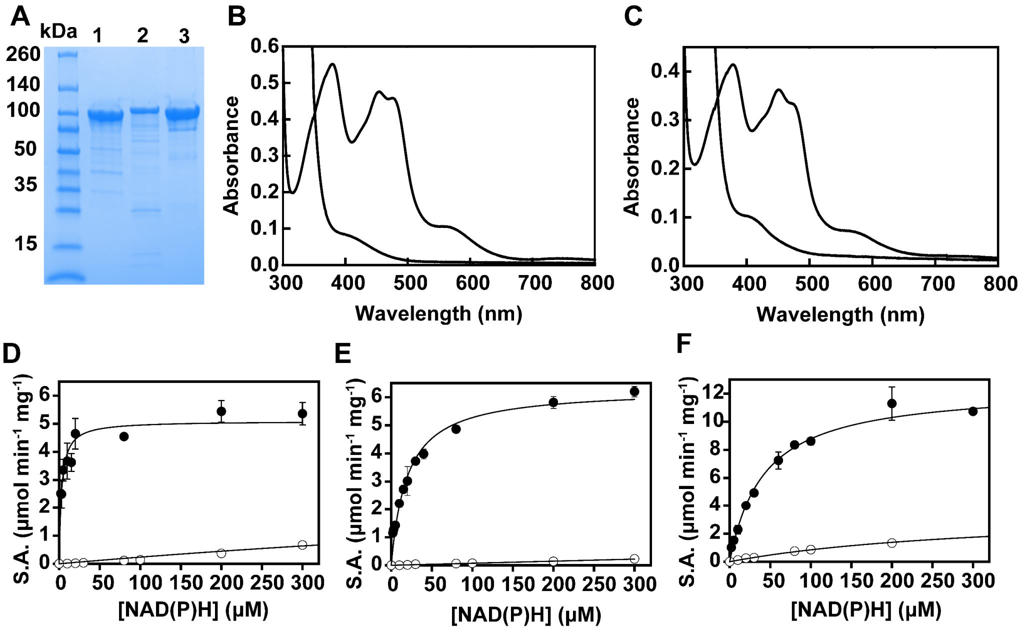


***Supplementary Figure S4. Initial purification of TvFDPF1-3 and their spectroscopic and kinetic analysis.*** **(a)** Purified *Tv*FDPF1 (lane 1), *Tv*FDPF2 (lane 2) and *Tv*FDPF3 (lane 3) (7 µg per lane). **(b-c)** UV-visible spectra of purified *Tv*FDPF1 and 3. Proteins were in buffer E at 40 µM (calculated based on the molecular weight of a monomer) as purified and after addition of 3 mM of sodium dithionite under aerobic conditions (bleached absorbance). The UV-visible spectrum of *Tv*FDPF2 suffers from extensive protein aggregation (shown in Supplementary Figure S5). **(d-f)** Michaelis-Menten analysis of the oxidase activity of *Tv*FDPF1-3 as described under "Experimental Procedures" with NADH (filled circles) and NADPH (open circles). All kinetic parameters of *Tv*FDPF1-3 are summarized in Table 2.

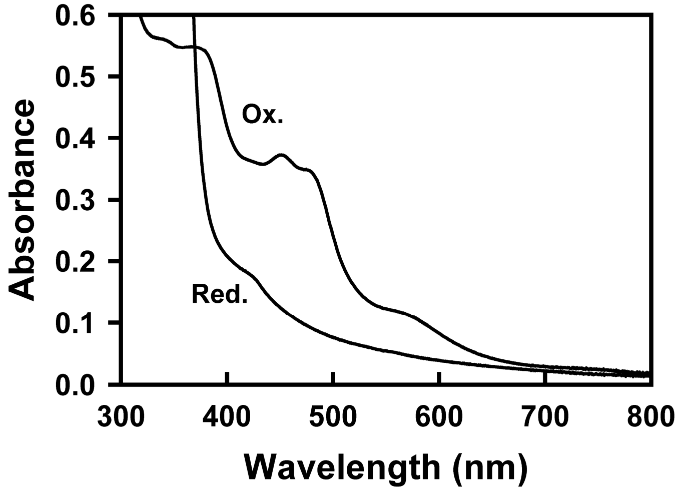


***Supplementary Figure S5. UV-visible spectra of purified TvFDPF2.*** Protein was in buffer E at 40 μM (calculated based on the molecular weight of a monomer) as purified and after addition of 3 mM of sodium dithionite under aerobic conditions. Aggregation of the protein can be seen by severe upward shift in the spectrum.

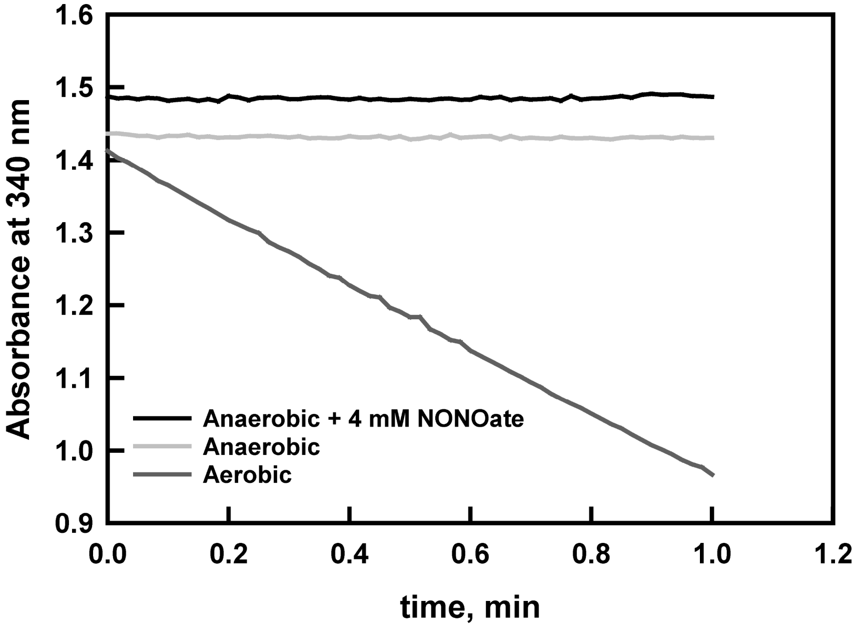


***Supplementary Figure S6.*** Activity with NO was quantified as described under "Experimental procedures". We estimated that the NOase activity of *Tv*FDPF3 was <2% of the NADH to $O_2$ specific activity.

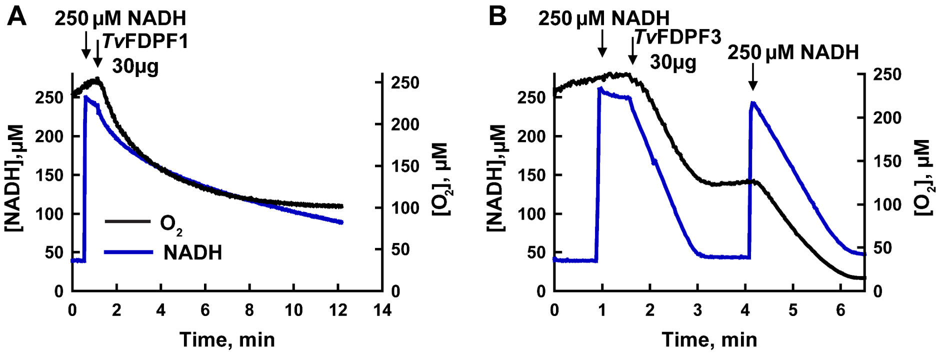


***Supplementary Figure S7. Kinetic characterization of the reaction with oxygen.*** Simultaneous measurements of oxygen (black traces) and NADH (blue traces) consumption by *Tv*FDPF1 (**a**) and *Tv*FDPF3 (**b**) that were performed as described under "Experimental procedures". Additions of NADH and enzymes are indicated with arrows.

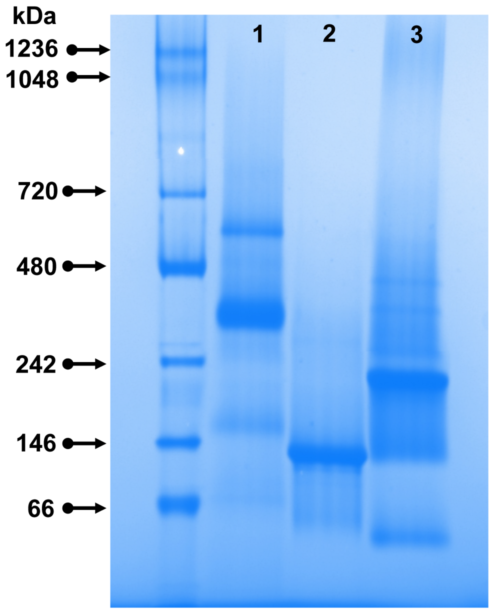


***Supplementary Figure S8.*** BN-PAGE of holo*Tv*FDPF3 (lane 1), *Lb*NOX (lane 2) and *Gi*NOX (lane 3). Based on the analytical gel-filtration the apparent molecular weight of *Lb*NOX is 195 ± 3 kDa and of *Gi*NOX is 241 ± 1 kDa. Note both *Lb*NOX and *Gi*NOX migrated on BN-PAGE accordingly to their expected molecular weights as determined by gel-filtration. The apparent molecular weight of holo*Tv*FDPF3 based on BN-PAGE is 322 ± 24 kDa based on (n = 3) independent experiments ± s.d.

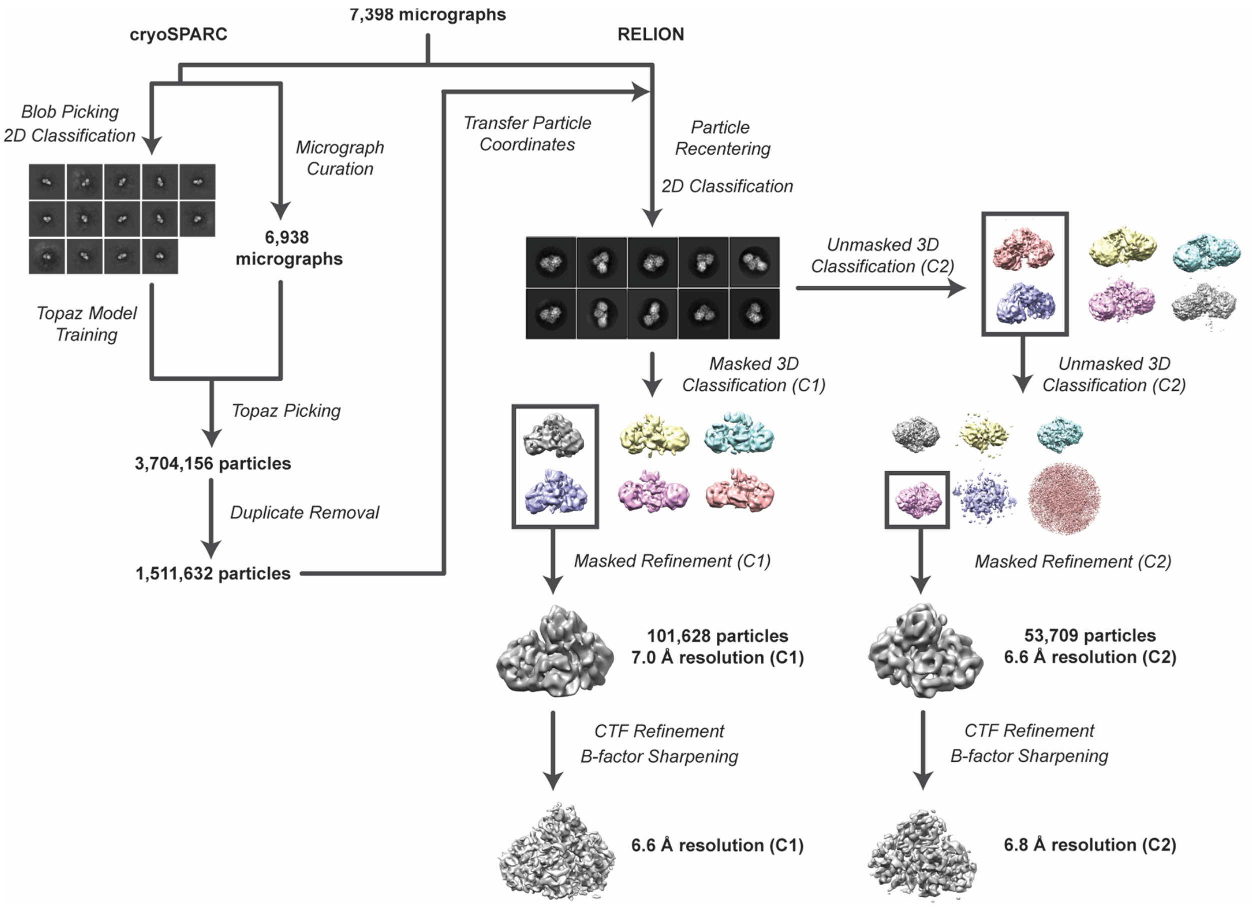


***Supplementary Figure S9. Simplified cryo-EM data processing workflow.***

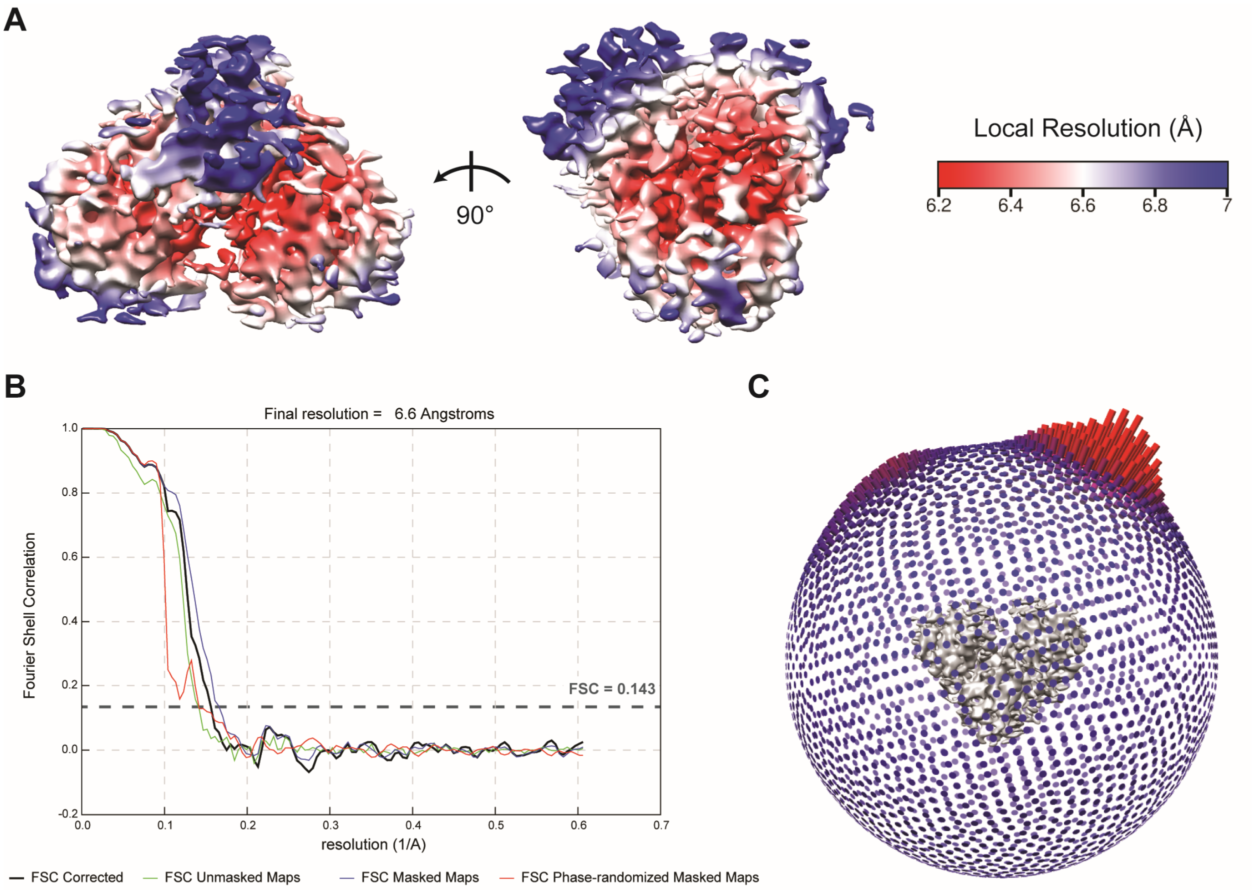


***Supplementary Figure S10. Cryo-EM map validation (C1 map).*** **(a)** Local resolution profile for the masked and sharpened C1 (no symmetry applied) holo*Tv*FDPF3 map. **(b)** Fourier shell correlation for the unmasked, masked, and phase-randomized masked C1 maps. The final corrected FSC is indicated in black. **(c)** Angular distribution of particle views in the final refined holo*Tv*FDPF3 C1 map.

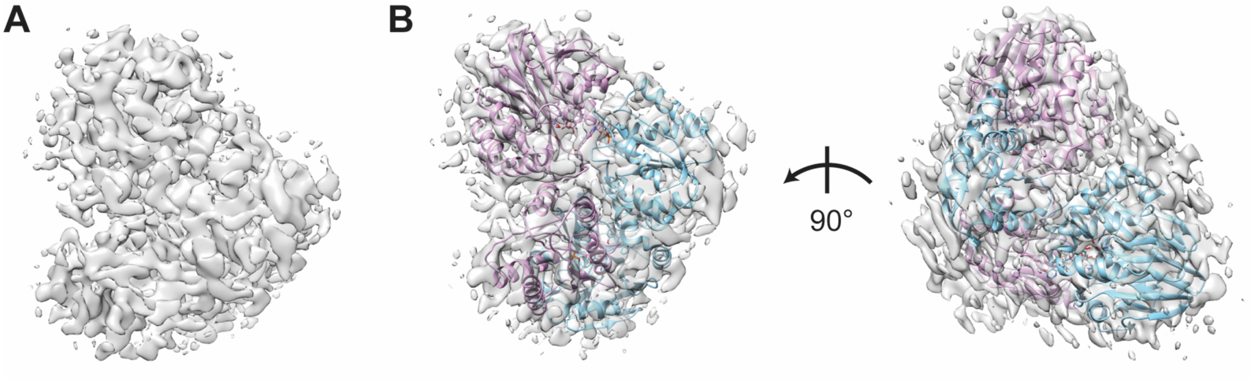


***Supplementary Figure S11. Homology model docking in C1 map.*** **(a)** C1-symmetric (no symmetry applied) electron density map for the holo*Tv*FDPF3 particles. **(b)** Homology models for the large and small subdomains of the holo*Tv*FDPF3 FDP-like domain docked into the C1-symmetric density map. The placement of the models indicates apparent C2 symmetry.

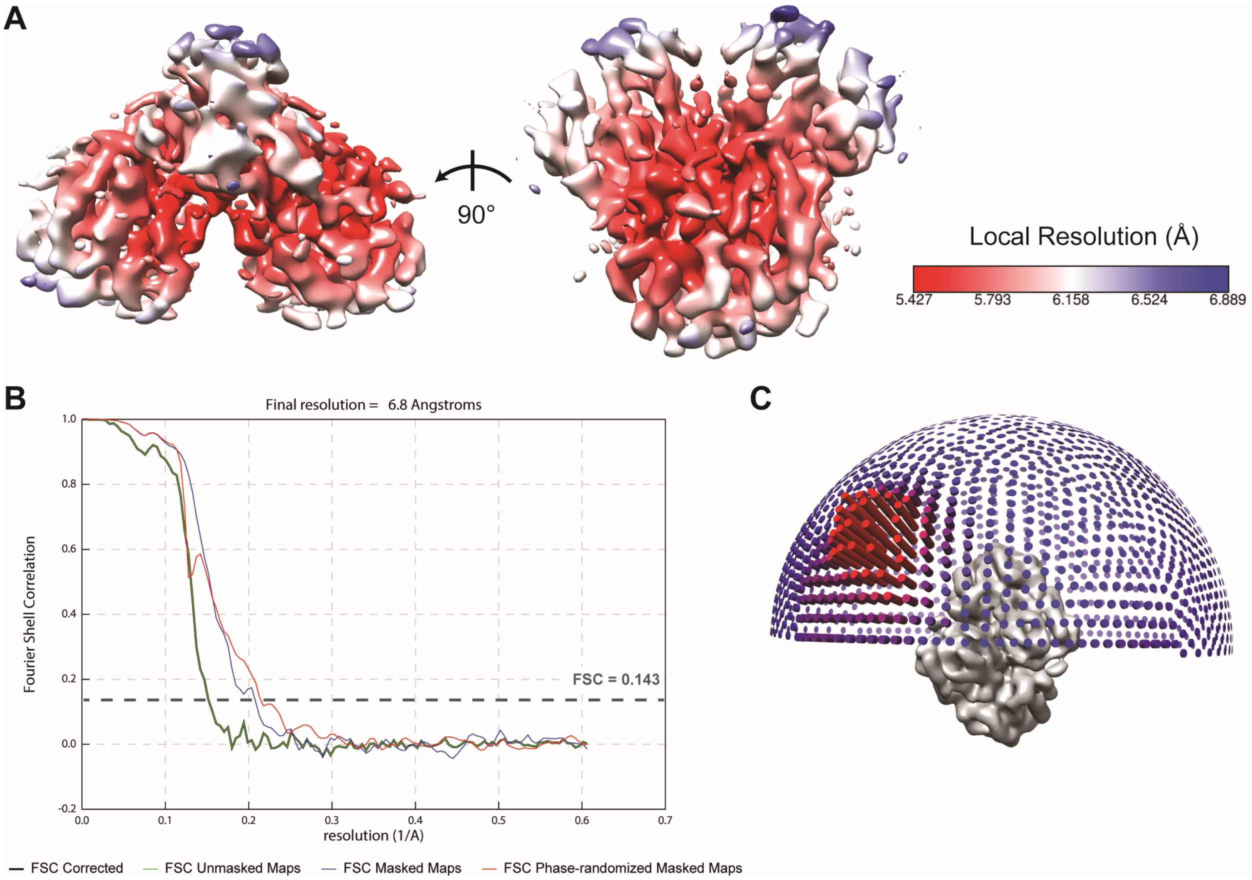


***Supplementary Figure S12. Cryo-EM Map Validation (C2 Map).*** **(a)** Local resolution profile for the masked and sharpened C2-symmetric holo*Tv*FDPF3 map. **(b)** Fourier shell correlation for the unmasked, masked, and phase-randomized masked C2 maps. The final corrected FSC is indicated in black. **(c)** Angular distribution of particle views in the final refined holo*Tv*FDPF3 C2 map.

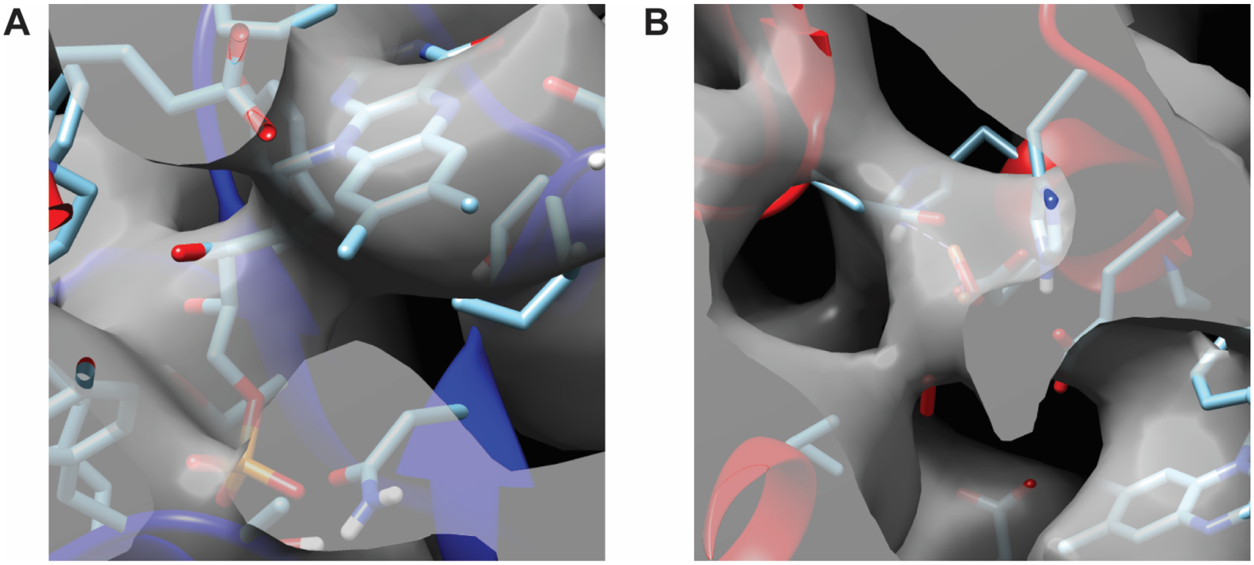


***Supplementary Figure S13. Ligand occupancy in electron density.*** Close-up view of the **(a)** FMN and **(b)** diiron ligands in the C2-symmetric electron density map.

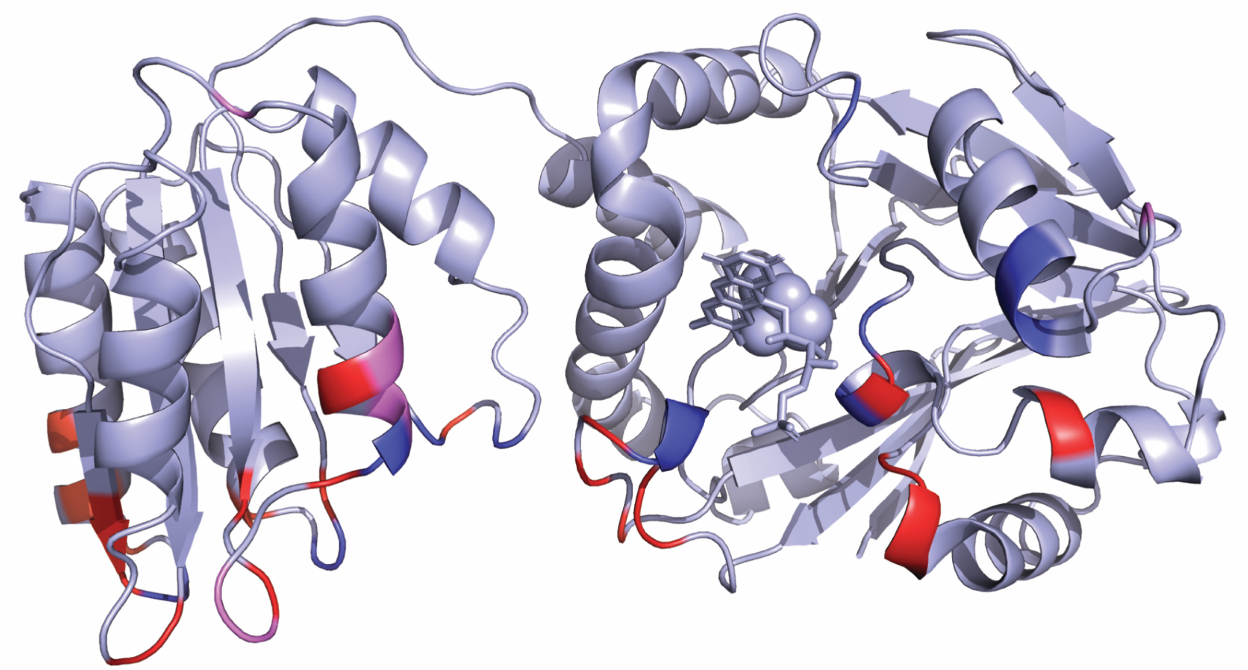

| Present in FDPF3 and ROO | | Unique to FDPF3 | | | Unique to ROO |
|---|---|---|---|---|---|
| D25 | Y268 | A22 | P85 | W334 | P120 |
| I26 | N297 | V23 | K92 | G357 | N326 |
| E84 | N300 | V27 | S266 | W358 | G327 |
| D86 | P331 | E29 | Y270 | I383 | P332 |
| D115 | W358 | K31 | I298 | K384 | D335 |
| I116 | F385 | K54 | N323 | E391 | H345 |
| W149 | R386 | E55 | I325 | K394 | |
| A267 | | R56 | V330 | | |

***Supplementary Figure S14. Inter-subunit contacts between holoTvFDPF3 protomers.*** Homology model of one subunit of the holo*Tv*FDPF3 FDP-like domain colored to identify interfacial residues (defined as residues in one subunit within 3.5 Å of any residue in the neighboring subunit). Residues colored blue form inter-protomer contacts in both the holo*Tv*FDPF3 model and the published structure of *D. gigas* ROO. Residues colored red form inter-protomer contacts unique to holo*Tv*FDPF3. Residues colored in purple form inter-protomer contacts in *D. gigas* ROO, but not in the model of holo*Tv*FDPF3. Residues are listed as numbered in holo*Tv*FDPF3.
